# Mutualists construct the ecological conditions that trigger the transition from parasitism

**DOI:** 10.1101/2021.08.18.456759

**Authors:** Léo Ledru, Jimmy Garnier, Matthias Rohr, Camille Noûs, Sébastien Ibanez

**Affiliations:** Univ. Grenoble Alpes, Univ. Savoie Mont Blanc, CNRS, LECA, 38000 Grenoble, France; CNRS, Univ. Grenoble Alpes, Univ. Savoie Mont Blanc, LAMA, 73000 Chambery, France; Laboratory Cogitamus

**Keywords:** mutualism, major transition, spatial structure, dispersal evolution, eco-evolutionary feedbacks, niche construction

## Abstract

The evolution of mutualism between hosts and initially parasitic symbionts represents a major transition in evolution. Although vertical transmission of symbionts during host reproduction and partner control both favour the stability of mutualism, these mechanisms require specifically evolved features that may be absent in the first place. Therefore, the first steps of the transition from parasitism to mutualism may suffer from the cost of mutualism at the organismic level. We hypothesize that spatial structure can lead to the formation of higher selection levels favouring mutualism. This resembles the evolution of altruism, with the additional requirement that the offspring of mutualistic hosts and symbionts must co-occur often enough. Using a spatially explicit agent-based model we demonstrate that, starting from a parasitic system with global dispersal, the joint evolution of mutualistic effort and local dispersal of hosts and symbionts leads to a stable coexistence between parasites and mutualists. The evolution of local dispersal mimics vertical transmission and triggers the formation of mutualistic clusters, counteracting the organismic selection level of parasites that maintain global dispersal. The transition occurs when mutualistic symbionts increase the density of hosts, which strengthens competition between hosts and disfavours hosts inhabiting areas dominated by parasitic symbionts: mutualists construct the ecological conditions that allow their own spread. Therefore, the transition to mutualism may come from an eco-evolutionary feedback loop involving spatially structured population dynamics.

## Introduction

Major transitions such as the evolution of chromosomes, eukaryotic cells, multicellularity and social groups have played a decisive role in the history of life. Most of these transitions resulted from the formation of a larger entity from smaller entities, smaller entities specializing within the larger ones, leading to transitions in individuality (Szathmáry and Smith, 1995). During transitions in individuality, smaller entities may either be like and unlike, resulting to a dichotomy between fraternal transitions arising from a division of labour among closely related units (such as multicellularity) and egalitarian transitions, where phylogenetically distant units come together to complement their functions in a larger unit Queller (1997); Szathmáry (2015). Egalitarian transitions are generally achieved through mutualistic symbiosis^1^ between a relatively large host and its symbiont (Bronstein, 2015; Drew et al., 2021) and constitute one of the main sources of new lineages, underlying the origin of the eukaryotic cell and photosynthetic eukaryotes for instance (Margulis and Sagan, 2002). In many cases symbionts are unicellular microbes which are hosted by large eukaryotes, the whole corresponding to a holobiont (Gilbert et al., 2012; Bordenstein and Theis, 2015); in other cases symbionts are multicellular organisms physically associated with their host at various degrees (e.g. plant-fungi, plant-ant, plant-seed eating pollinator). While symbionts depend on their host from the start, hosts often become dependent on the symbionts during later stages (Roughgarden, 1975), e.g. for reproduction or resource acquisition, eventually making the transition irreversible.

For a transition to occur and persist, evolutionary conflicts between the subentities must not overtake the whole’s fate. In the case of fraternal transitions, this is prevented by the strong relatedness between subentities (Hamilton, 1964b,a; Queller, 2000; Fisher et al., 2013). However, in the case of egalitarian transitions, the subentities generally belong to different species. Thus, it can be advantageous for them to remain autonomous and exploit the other subentities. This parasitic behaviour occurs at the expense of the whole, as for the tragedy of the commons (Garrett, 1968; Hardin, 1998). For instance, a symbiont may remain parasitic rather than collaborate with its host (Drew et al., 2021). The resulting evolutionary conflict might be circumvented by vertical transmission of the symbionts, which ensures that all subentities share a common fate (Wilson and Sober, 1994). As a result, vertical transmission of symbionts indeed promotes the transition to mutualism (Smith, 1998; Herre et al., 1999; Wilkinson and Sherratt, 2001; Ferdy and Godelle, 2005; Kerr and Nahum, 2011; Akçay, 2015; Estrela et al., 2016; Queller and Strassmann, 2016; Doebeli and Knowlton, 1998), although symbionts vertically transmitted can persist without becoming mutualists (Saikkonen et al., 2002).

The importance of vertical transmission has been highlighted by experiments on microbial systems (Sachs et al., 2011; Shapiro et al., 2016; King et al., 2016; Shapiro and Turner, 2018) as well as *in natura* observations of a Wolbachia-insect system (Weeks et al., 2007). However, in many mutualistic systems, the symbiont is transmitted horizontally (Wilkinson and Sherratt, 2001), such as legume-rhizobium (Denison and Kiers, 2004), squid-vibrio (McFall-Ngai, 2014), mycorrhizae (Allen, 1991), endophytes (Saikkonen et al., 2004) or plant-ants (Bronstein et al., 2006; Rico-Gray and Oliveira, 2008). In such cases, several mechanisms such as partner choice, sanction or fidelity can counteract the selection for selfishness (Genkai-Kato and Yamamura, 1999; Wilkinson and Sherratt, 2001; Sachs et al., 2004; Foster and Wenseleers, 2006; Estrela et al., 2016; Akçay, 2017; Sachs et al., 2010b). For instance, in legume-rhizobium, mycorrhizal and plant-ant associations, the plants can sanction the less beneficial (or even detrimental) symbionts by allocating them fewer resources (West et al., 2002; Kiers et al., 2003; Denison and Kiers, 2004; Edwards et al., 2006; Bever et al., 2009; Akçay, 2015). However, it is unclear whether these mechanisms are present at the beginning of the transition to mutualism. Since they require the evolution of complex and specific traits, they may occur in later stages, providing additional stability to the system. In the absence of such traits, what mechanism could promote the transition in the first place? Using a theoretical model, the present work aims to show that the joint evolution between mutualistic effort and local dispersal of hosts and symbionts leads to a positive association between mutualistic hosts and symbionts and subsequently triggers the formation of mutualistic clusters.

A similar issue exists with respect to the evolution of altruism^2^, since partner choice and control mechanisms, such as voluntary reciprocal altruism (Axelrod, 1981), may be restricted to higher animals or may appear during later evolutionary stages. In line with the intuition of Darwin (1871), spatial structure has been recognized as a general mechanism promoting the transition to altruism (Mitteldorf and Wilson, 2000; Lion and Van Baalen, 2007, 2008; Débarre et al., 2012). Spatial structure generally stems from local dispersal, which triggers the formation of clusters dominated by altruistic organisms, while organisms with similar phenotypes are positively assorted in space (Wilson and Dugatkin, 1997; Pepper, 2007). The balance between organismic-level selection favouring cheaters and cluster-level selection favouring altruists ultimately determines the evolutionary outcome (Van Baalen and Rand, 1998; Mitteldorf and Wilson, 2000). Moreover, the joint evolution of cooperation and dispersal can allow the emergence of altruism, with spatial clusters of altruistic organisms promoting the persistence and spread of altruistic phenotypes (Koella, 2000; Le Galliard et al., 2005; Hochberg et al., 2008; Purcell et al., 2012; Mullon et al., 2018). Empirical evidence on the evolution of reduced virulence (Boots and Mealor, 2007; Szilágyi et al., 2009), the evolution of altruism (Harcombe, 2010; Eldakar et al., 2010), and the evolution of restraint predation (Kerr et al., 2006) also supports the crucial role of the spatial structure.

Similarly, spatial structure can allow mutualists to overcome non-mutualists (Yamamura et al., 2004; Akçay, 2017; Doebeli and Knowlton, 1998; Frank, 1994), and this can come along with the evolution of dispersal (Mack, 2012). However, this may not be sufficient to account for the transition from parasitism to mutualism, since parasitic symbionts should discourage hosts from initiating the transition, whereas non-mutualists have a weaker impact (Yamamura et al., 2004; Mack, 2012). In the case of holobionts, starting from free living bacteria, Sachs et al. (2011) documented 27 transitions towards parasitism, 9 directly towards mutualism and 3 towards commensalism, whereas the transition from parasitism to mutualism occurred only 3 times. This highlights that the transition from parasitism to mutualism, although feasible, is relatively infrequent, and calls for a theoretical understanding of the mechanisms involved. Moreover, in previous attempts mutualistic efforts were initially polymorphic but were not subject to mutations (Mack, 2012). In that case, mutualistic clusters cannot be invaded from inside through parasitic mutations, which favours mutualism. The present work therefore constitutes, to our knowledge, the first spatially explicit eco-evolutionary model where the mutualistic efforts and dispersal abilities of hosts and symbionts coevolve, beginning from a parasitic interaction. If some hosts and symbionts simultaneously become mutualists and start dispersing locally, this may lead to the formation of mutualistic host-symbiont clusters producing more offspring than in areas where hosts are mainly associated with parasitic symbionts, thereby initiating the transition. Meanwhile, parasitic symbionts should continue dispersing globally and invade the mutualistic clusters, which could homogenize the spatial structure and compromise the transition. Also, densely populated mutualistic clusters might suffer from intraspecific competition between hosts, unless competition acts on a large spatial scale. In sum, it is unclear whether mutualists will invade, whether mutualists will replace parasites, or whether both strategies will coexist, as is often the case in nature (e.g. Després and Jaeger, 1999; Borges, 2015; Saikkonen et al., 2004).

The concept of major transitions also implies that the host and the symbiont become dependent upon each other (Szathmáry and Smith, 1995; Szathmáry, 2015; Nguyen and Baalen, 2020), with each partner needing the other to perform essential functions like nutrient provisioning (Fisher et al., 2017). Dependence is often accompanied by gene loss and gene exchange, rendering the transition irreversible (Estrela et al., 2016). Most symbionts cannot live freely and therefore completely depend on their host, but most hosts can complete their life cycle without their symbiont (e.g., in plant-ant, plant-fungi or legume-rhizobium mutualisms) and several reverse pathways are possible from mutualism to parasitism (Sachs and Simms, 2006; Werner et al., 2018; Week and Nuismer, 2021). However in some cases hosts depend on their symbiont, for instance the intracellular bacterial symbiont *Buchnera aphidicola* provides essential amino acids to its aphid host (Akman Gündüz and Douglas, 2009; Bennett and Moran, 2015). Since the present work focuses on the transition and not on later stages, we will not assume that hosts depend on their symbionts for their physiology or development, which would render the transition irreversible *by construction*. Instead, hosts will always be able to produce offspring when alone. Nevertheless, the number of offspring produced by the hosts will depend on the mutualistic efforts of both species as well as on the population densities, which are expected to change during the transition. Under these altered ecological conditions, isolated hosts may exhibit a negative population growth rate, although they are physiologically able to produce offspring. This mechanism is hereafter called *ecological dependence*.

To sum up, we will tackle the following issues:

- **Main hypothesis:** In the absence of vertical transmission and partner control, we expect that the transition from parasitism to mutualism can occur when the mutualistic efforts of both hosts and symbionts jointly evolve with local dispersal.
- **H1:** The formation of mutualistic clusters should be necessary for the initiation of the transition. The emergence of spatial structure should come along with the transition.
- **H2:** By maintaining global dispersal, non-mutualistic hosts and parasitic symbionts should be able to coexist with mutualists.
- **H3:** The transition to mutualism is due to the relatively higher fecundity of mutualistic clusters.
- **H4:** If competition between hosts is mostly local, this should hamper the formation of mutualistic clusters, thereby preventing the transition.
- **H5:** We expect that mutualistic hosts will become ecologically dependent on their symbiont.

To investigate these hypothesis, we built an agent-based model using a two-dimensional space lattice that supports an autonomous host and a host-dependent symbiont. Hosts compete for space and other resources, while symbionts compete for available hosts. This situation occurs in many biological systems, such as plant-fungi, plant-seed eating pollinator, plant-ant, and multicellular eukaryotes hosting bacteria. Less intimate associations like cleaning mutualisms or plant-pollinator interactions may also fit, provided that the animal is specialized and dependent on its host. To model the transition from parasitism, the symbiont is initially detrimental to the host, and the host provides it the minimal energy possible without any spontaneous mutualistic effort, as would be the case after an antagonistic evolutionary arms race. Moreover, the host-parasite system is ecologically viable even in the absence of any mutualistic agent in the landscape. At first, both species disperse globally; this situation corresponds to the most disadvantageous conditions for the emergence of mutualism. Through continuous mutations, mutualistic and locally dispersing symbionts and hosts can appear. The mutualistic effort encompass the provision of resources, shelter, immunity, anti-predator behaviours, digestive enzymes or any other type of benefit provided that this occurs at some cost. If mutualistic symbionts manage to persist for a while, they eventually change the population dynamics, triggering feedback on their own evolutionary dynamics. In addition to these general hypotheses, no assumptions specific to a particular biological system were required.

## Methods

### Main rules of the model

Our model considers two types of agents, hosts and symbionts, living on the same two-dimensional space lattice. The interaction between the two species occurs when they share the same cell. Each cell can assume three states: i) empty, ii) occupied by a solitary host, with only one host per cell), iii) occupied by a host-symbiont couple, with only one symbiont per host (but see Appendix A.7 for a relaxation of this assumption). Each organism bears two traits, an interaction trait *α* and a dispersal trait *ε*, which both influence fecundity. At every time point, agents undergo the following steps (see appendix A.1 and Figure A1 for more details):

- The host and symbiont die with fixed probability *m*.
- They produce offspring. The average offspring number of a parent depends on its traits and on its interactions with their cell-sharing partner, if any.
- The offspring are dispersed according to the parental trait *ε*. For instance, the dispersal abilities of akenes depend on the parental genotype.
- The host offspring may establish only in empty cells, while the symbiont offspring can only establish in cells already occupied by a solitary host. If several organisms come to implant in the same cell, a uniform lottery determines which one will implant, while the others die.
- The offspring traits mutate with a given probability, which will affect their own interaction with hosts/symbionts and the dispersal of their future offspring. In nature mutations occur as soon as offspring are produced, instead in the model only the surviving offspring mutate, which saves computation time.

### Fecundity and mutualism/parasitism

Each agent produces offspring according to a Poisson distribution with parameter *f*, which corresponds to its fecundity. The fecundity defines the average number of offspring per agent. It results from an interaction fecundity positively dependent on the trait of its cell-sharing partner and a mutualistic cost negatively dependent on its trait.

Specifically, the fecundity of a symbiont *f*^*s*^ of trait *α*_*s*_ in interaction with a host of trait *α*_*h*_ is defined by:

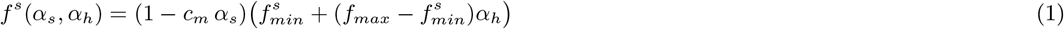

where *c*_*m*_ is the maximal mutualistic cost and 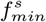 and *f*_*max*_ are the minimal and maximal interaction fecundity of symbionts. Similarly, the fecundity of a host *f*^*h*^ of trait *α*_*h*_ in interaction with a symbiont of trait *α*_*s*_ is defined by:

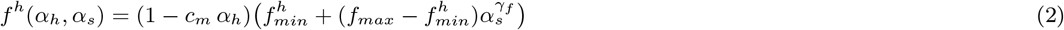

where 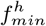 is the minimal interaction fecundity of hosts. The parameter *γ*_*f*_ describes how fast the fecundity of the host increases with the interaction trait of the symbiont *α*_*s*_. Since we are interested in the emergence of mutualism, the parameter *γ*_*f*_ describes the mutualistic strength of the symbiont on the host fecundity. In our model, we set the mutualistic strength of the host on the symbiont fecundity to *γ*_*s*_ = 1.

Since hosts are autonomous, in absence of symbionts, their fecundity *f*^*ha*^ only depends on their trait *α*_*h*_:

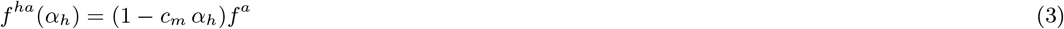

where the fecundity alone *f*^*a*^ ranges between the minimal and maximal interaction fecundity: 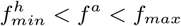. As a result, the establishment of a symbiont with a low interaction trait 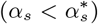 reduces the fecundity of the host; the symbiont is parasitic. Instead, a symbiont with a large interaction trait 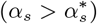 enhances the host’s fecundity; the symbiont is mutualistic. The threshold 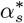 is defined by 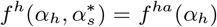 (see appendix A.1 for mathematical derivation of the threshold). In the simulations, 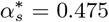 (Figure 1).

**Figure 1:**
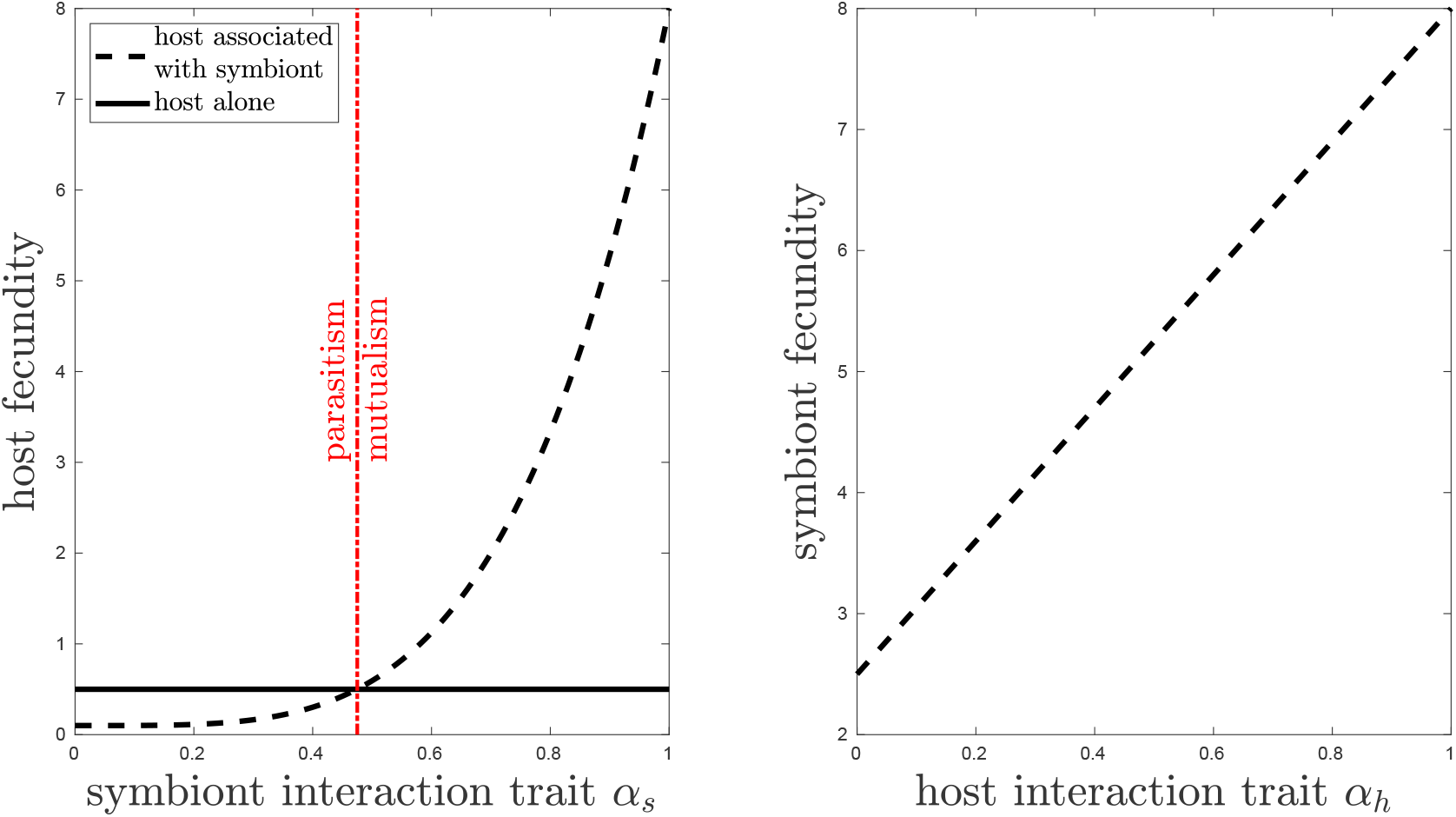
Fecundity of hosts *f*^*h*^ and symbionts *f*^*s*^ according to the interaction trait of their partners (dashed black lines). Plain black line corresponds to the fecundity of a solitary host *f*^*a*^. The dashed red line in panels (a) and (b) corresponds to the threshold 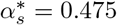 separates symbionts, which either reduce or enhance the fecundity or their host (appendix A.1).

### Mutation

Offspring inherit traits from their parents with variability due to mutations. The effects of mutations on each trait are independent. However, the distribution of mutation effects does depend on the trait of the parents. We use a Beta distribution with shape parameters (1, 3) to describe the amplitude of these effects, which could be either beneficial or detrimental. This mutation kernel allows for rare mutations with large effects. However, these effects can not exceed a maximal mutation size set to *β*_*max*_ = 0.5 (see Figure A3 in appendix A.1 for details).

### Dispersal

The parents do not disperse, while their descendants disperse either locally in one of the 8 cells around the parent or globally, with a uniform distribution across the entire space (see Figure A5 for a sketch of the process). The dispersal trait *ε* is defined as the proportion of offspring dispersed globally, as in Kéfi et al. (2007, 2008). These two modes of dispersal correspond to a mixture of short and long distance dispersal events. For instance, fleshy fruits may be dispersed either by small birds having a short-distance behaviour, or by mammals and large birds which disperse the seeds at long distances (Jordano et al., 2007). Fruits may also remain unconsumed and fall locally. Depending on the fruit’s traits, its propensity to be consumed by either type of frugivores may vary among organisms, which is captured by the dispersal trait *ε*. Since the investment in global dispersal may reduce fecundity (Harada, 1999; Bonte et al., 2012), we assumed a linear trade-off between fecundity and dispersal: *f*_*e*_ = (1 − *d ε*) *f*, with *f*_*e*_ the effective fecundity and *d* the dispersal cost intensity, which is the same for both hosts and symbionts.

### Competition

Hosts compete for empty cells, especially if they disperse locally. Beside space, hosts may also compete with each other for resources like water, light or food. In order to test hypothesis H3 we introduced intraspecific density-dependent competition, acting either at the local or the global scale. For instance, competition for light only involves the closest neighbors while competition for the water table might act at the entire space scale. The competition scale parameter *w*_*h*_, ranging in [0, 1], weights the effect of the local density 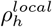 and the global density 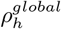 of host on the competition. Competition reduces the establishment probability *P*_*I*_ of the offspring:

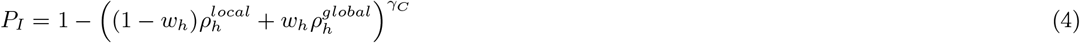

The local host density 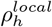 corresponds to the host density in the 8 neighbouring cells surrounding the offspring, while the global density 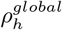 corresponds to the host density over the entire landscape (see Figure A2 for a schematic representation). The parameter *γ*_*C*_ corresponds to the inverse of the competition strength. Indeed, the establishment probability is increasing with respect to *γ*_*C*_. Thus, the competition is strong when *γ*_*C*_ *<* 1 (sub-linear function), while it is weak when *γ*_*C*_ ≥ 1 (super-linear function).

### Parasitic system and transition

To tackle the issue of transition to mutualism, we assume that the system is viable without mutualism (see appendix A.2 for details). More precisely, in the absence of mutation, the per capita growth rate at low densities of hosts is *F*_0_ = (1 − *m*)(1 + *f*^*a*^). In our study, we have chosen parameters (see Table 1) for which *F*_0_ is greater than 1 so that if we start with a large density of hosts initially, the probability of extinction is 0. Moreover, under our parameter ranges, the system stabilizes around a demographic equilibrium called the “parasitic system” where host density is around 0.15 and the symbiont density is around 0.1 (see Figure 2b-c and appendix A.2 for mathematical details on the stability of an approximation model). From our perspective, this situation is the worst-case scenario because interactions are parasitic and dispersal cost is minimal. Then, mutualistic symbionts can appear by mutation, which generates approximately 2% of mutualistic symbionts in the population (see dashed purple curve in Figure 2c). Natural selection eventually leads to a significant increase of the percentage of mutualistic symbionts, far above the 2% generated by mutations (Figure 2). In the simulations, a high density of mutualistic symbionts indeed persists in the long term when the percentage of mutualistic symbionts stands above 10% (Figure A8), which therefore characterizes the transition to mutualism. The transition time was defined as the time at which the percentage of mutualistic symbionts rises above this threshold.

**Table 1:** List of parameters and their reference values used for the simulations. The parameters of host and symbiont fecundities are determined to ensure the viability of the antagonistic system, therefore they are fixed because they are constitutive of the model.

| Parameters |  | Reference values | Sensitivity analysis range |
| --- | --- | --- | --- |
| $m$ | probability of mortality | 0.06 | [0.005; 0.15] |
| $c_m$ | maximum mutualistic cost | 0.3 | [0; 1] |
| $f_{max}$ | maximal host and symbiont interaction fecundity | 8 | fixed |
| $f_{min}^h$ | minimal host interaction fecundity | 0.1 | fixed |
| $f_{min}^s$ | minimal symbiont interaction fecundity | 2.5 | fixed |
| $f^a$ | maximal solitary host fecundity | 0.5 | fixed |
| $\gamma_f$ | mutualistic strength on the host fecundity | 4 | fixed |
| $\beta_{max}$ | maximum mutation size | 0.5 | [0.1; 1] |
| $w_h$ | scale of host competition | 1 | 0 or 1 |
| $\gamma_c$ | inverse of host competition strength | 0.2 | [0.1; 2] |
| $d$ | dispersal cost | 0 | [0; 1] |

**Figure 2:**
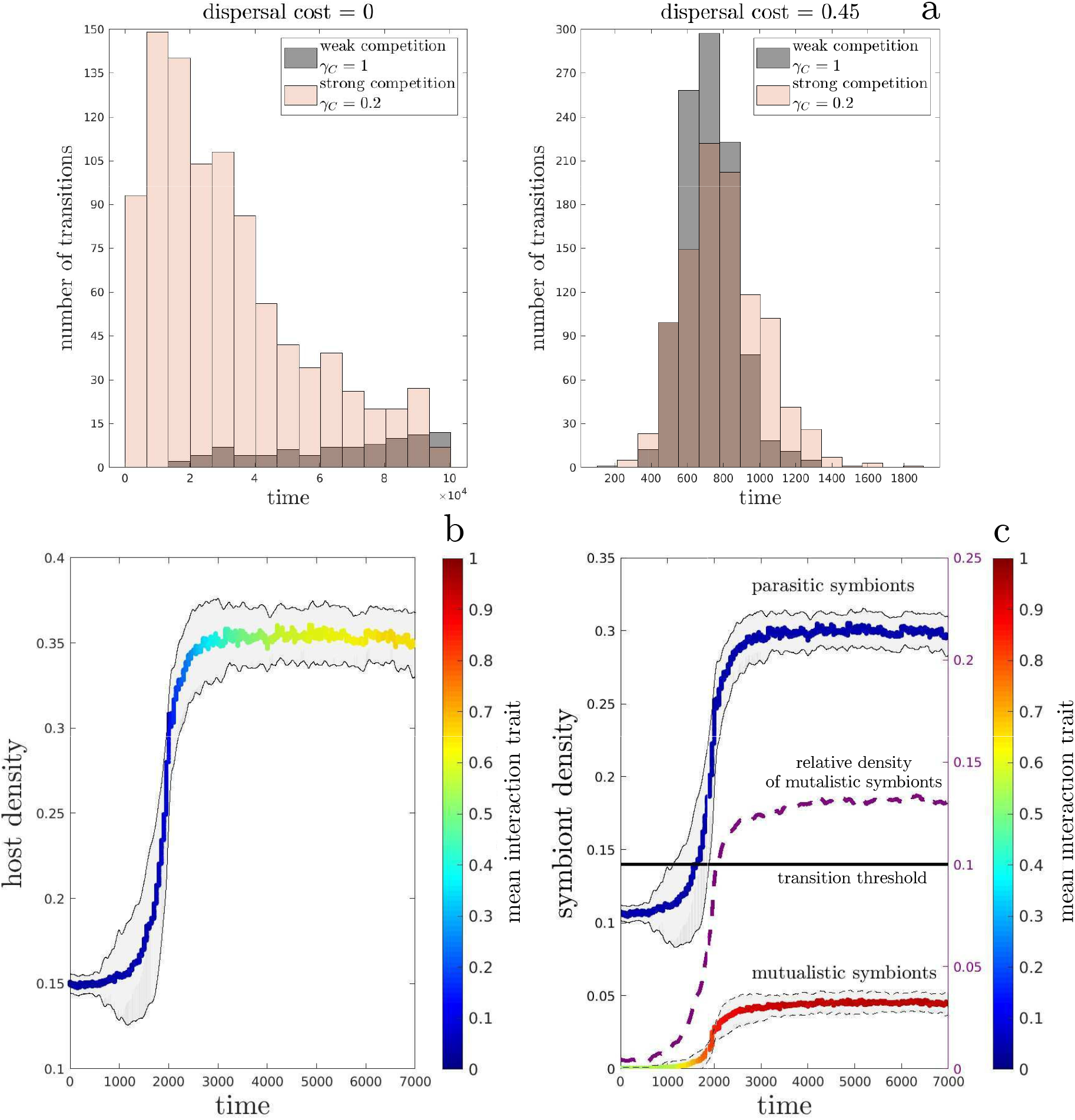
a) Histograms of the number of transitions over 1000 simulations as a function of time, with a maximum projection time of 10^5^. Without dispersal cost there are a total of 86 transitions when the competition is weak and 951 when the competition is strong. With dispersal cost there are 1000 transitions whether the competition is weak or strong. Panels b) and c) represent the host and symbiont densities over time averaged over 100 simulations (coloured plain curves) under strong competition *γ*_*C*_ = 0.2 and no dispersal cost *d* = 0 and with a maximum projection time of 10^4^ steps. The densities correspond to the proportion of occupied cells. The time series are adjusted so that all simulations have a transition time *t* = 2000. The colour gradient corresponds to the mean interaction trait *α*, and shaded regions correspond to the standard deviation for densities. In panel c), the purple dotted line and the right y-axis show the relative density of mutualistic symbionts, and the black line indicates the 10% transition threshold. For all panels, other parameters are 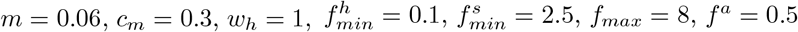 and *β*_*max*_ = 0.5

### Assortment and aggregation indices

To investigate the spatial structure, which comes along with the transition to mutualism, we compute assortment indices: intraspesific indices measuring the spatial autocorrelation among hosts and symbionts and an interspecific index quantifying the correlation between phenotypes of host and symbiont sharing the same location. More specifically, the intraspecific indices compute the similarity between the trait of an organism and the traits of its neighbors located in the 8 cells around it, and compare it with the similarity between the organismic trait and the mean trait over the landscape (details in appendix A.1). If the intraspecific index is positive (respectively negative), it means that on average the neighbors of any organism share similar (respectively dissimilar) traits. Similarly, the interspecific index is positive if hosts and symbionts sharing the same cell have similar interaction traits. Spatial aggregation indices for hosts, mutualistic symbionts and parasitic symbionts were also computed, measuring the formation of clusters (appendix A.1 for details).

## Results

In the following, the maximum cost of mutualism *c*_*m*_ is 30%, and the other parameters are set to satisfy the viability of the parasitic system (Table 1 in appendix A.1 and appendix A.2 for a discussion of the effect of the cost of mutualism).

### The transition from parasitism to mutualism

Our main objective was to investigate whether the transition to mutualism is possible starting from a viable parasitic system, without dispersal cost, which constitutes the most stringent condition for the transition. In that case, the transition is more likely to occur under strong (*γ*_*C*_ = 0.2) intraspecific host competition (with frequency 0.95) than under weak (*γ*_*C*_ = 1) competition (0.086). Moreover, when the transition succeeds, it occurs more rapidly under strong competition (median transition time around 2.5.10^4^) than under weak competition (median transition time around 7.10^4^, Figure 2a). When the cost of dispersal is large (*d* = 0.45) the transition occurs systematically (with frequency 1) and the median transition time is much lower (around 7.10^2^), regardless of the strength of competition (Figure 2a). Dispersal cost was therefore used as an instrumental tool to speed up the transition when necessary.

The transition begins with weakly mutualistic symbionts, which rapidly increase their mutualistic effort toward 1 (Figure 2c). In contrast, the increase of the average host interaction trait is delayed in response to the symbionts’ transition (Figure 2b). Moreover, the transition does not occur at the expense of parasitic symbionts; on the contrary their population density benefit from the increase in host density triggered by the mutualistic symbionts (Figure 2c).

Since the symbiont population is monomorphic at the beginning of every simulation, the two distinct phenotypic clusters visible in Figure 3a indicate that both traits diverged, resulting in two classes of symbionts: parasitic global dispersers (*α*_*s*_ *<<* 1 and *ε* ∼ 1) and mutualistic local dispersers (*α*_*s*_ ∼ 1 and *ε <<* 1). Furthermore, the mutualistic and dispersal traits of symbionts evolve at the same time, during the transition (details not shown). Conversely the host traits do not diverge; their joint evolution leads to a negative correlation between global dispersal and mutualism intensity (Figure 3b, R^2^=0.102). After the transition, most hosts provide a non-zero mutualistic effort to the symbiont (most *α*_*h*_ *>* 0.2).

**Figure 3:**
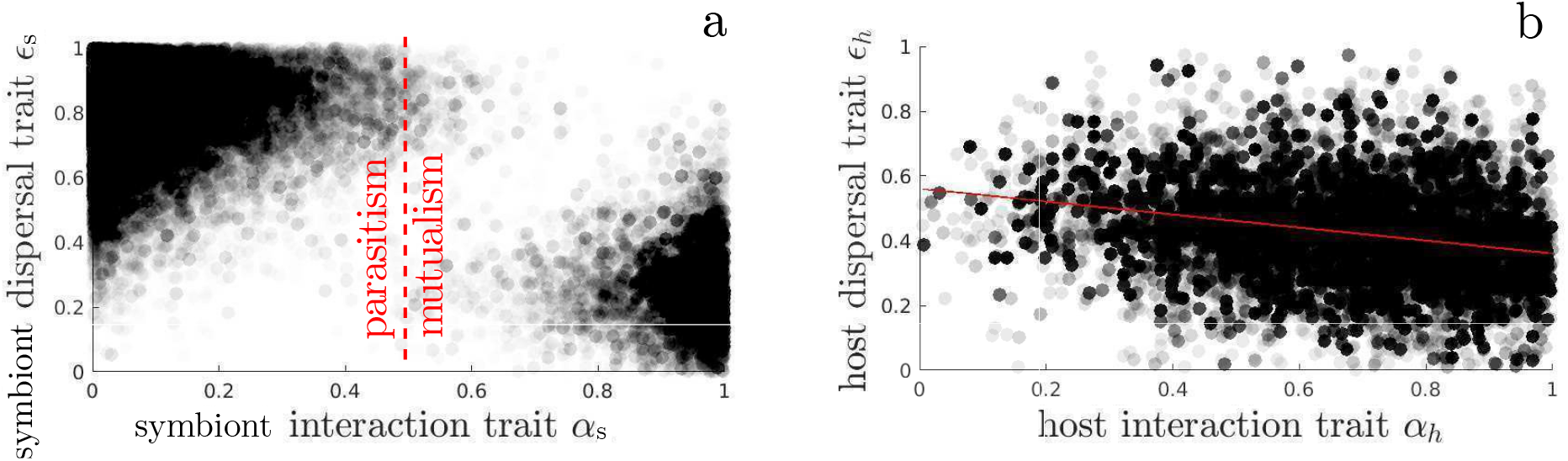
Post transition traits distribution (*ε, α*) of symbionts (panel a) and hosts (panel b). The dashed red line in panel a indicates the threshold 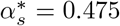 above which a symbiont benefits its host. The plain red line in panel b shows the linear regression between host traits (R^2^=0.102). Distributions corresponds to 100 simulations with strong competition *γ*_*C*_ = 0.2, no dispersal cost *d* = 0 and with a maximum projection time of 10^4^ steps. Other parameters are 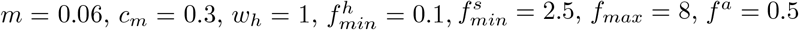 and *β*_*max*_ = 0.5.

The assortment indices indicate that after the transition to mutualism the organisms of both species are locally similar. Moreover, hosts and symbionts sharing the same location also tend to have the same interaction behaviour (Figure 4a). The intraspecific assortment is stronger than the interspecific assortment, which is not surprising since the formation of the intraspecific spatial structure simply requires a sufficient proportion of local dispersal. The aggregation indices (appendix A.1) behave similarly, after the transition the spatial aggregation of hosts, parasitic symbionts and mutualistic symbionts all increase, and the parasitic and the mutualistic symbionts reach the same level of aggregation (Figure A7). These results together indicate that the transition to mutualism comes along with the emergence of a spatial structure, with clusters of mutualistic hosts and symbionts (Figure 4c).

**Figure 4:**
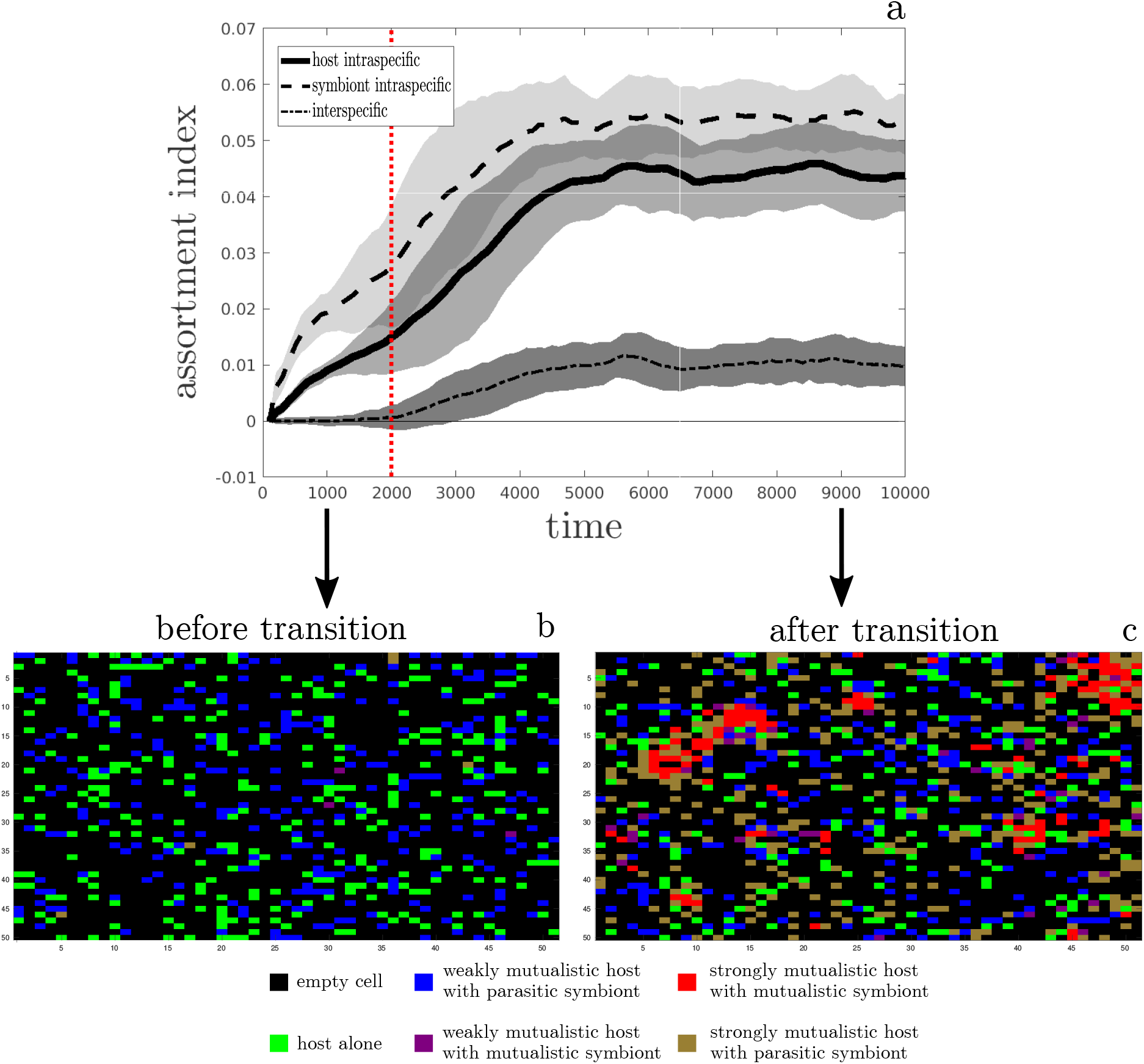
a) Spatial structures are described by the assortment index measuring the intraspecific assortment between hosts (plain line), the intraspecific assortment between symbionts (dashed line) and the interspecific assortment between hosts and symbionts (dash-dot line). Results are averaged over 100 simulations with strong competition *γ*_*C*_ = 0.2 and no dispersal cost *d* = 0. The time series are adjusted so that all simulations have a transition time *t* = 2000 (red dotted line). Grey areas show the standard deviation. The threshold separating mutualistic and antagonistic symbionts is as in Figures 1 and 3. b)-c) Snapshots of a region of 40×40 cells before (panel b) and after (panel c) the transition to mutualism. For the sake of the figure, a host is considered weakly mutualistic if its interaction trait is less than 0.5 and strongly mutualistic if it is greater than 0.5.Other parameters are 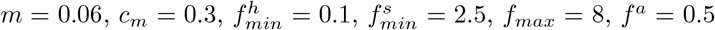 and *β*_*max*_ = 0.5

### The effect of competition between hosts

Figure 2a shows that the host competition promotes the transition to mutualism; we next investigate its quantitative effect on the percentage of mutualistic symbionts. The following results were obtained using a large dispersal cost (*d* = 0.45) to reduce the mean time of transition and thus save computational time.

The competition strength *γ*_*C*_ increases the percentage of mutualistic symbionts after the transition when competition is global, i.e. when hosts compete with all the hosts present in the landscape (Figure 5a). However, the transition can occur even in the absence of host competition, if the cost of mutualism is sufficiently low (e.g., a maximum cost of only 10% instead of 30% as in previous simulations, details not shown). When competition is more local the percentage of mutualistic symbionts decreases drastically, until it drops below the transition threshold (Figure 5b). In the absence of dispersal cost, when competition is reduced after the transition to mutualism, the system switches back to the parasitic state (Figure 5c, see Figure A9 for details).

**Figure 5:**
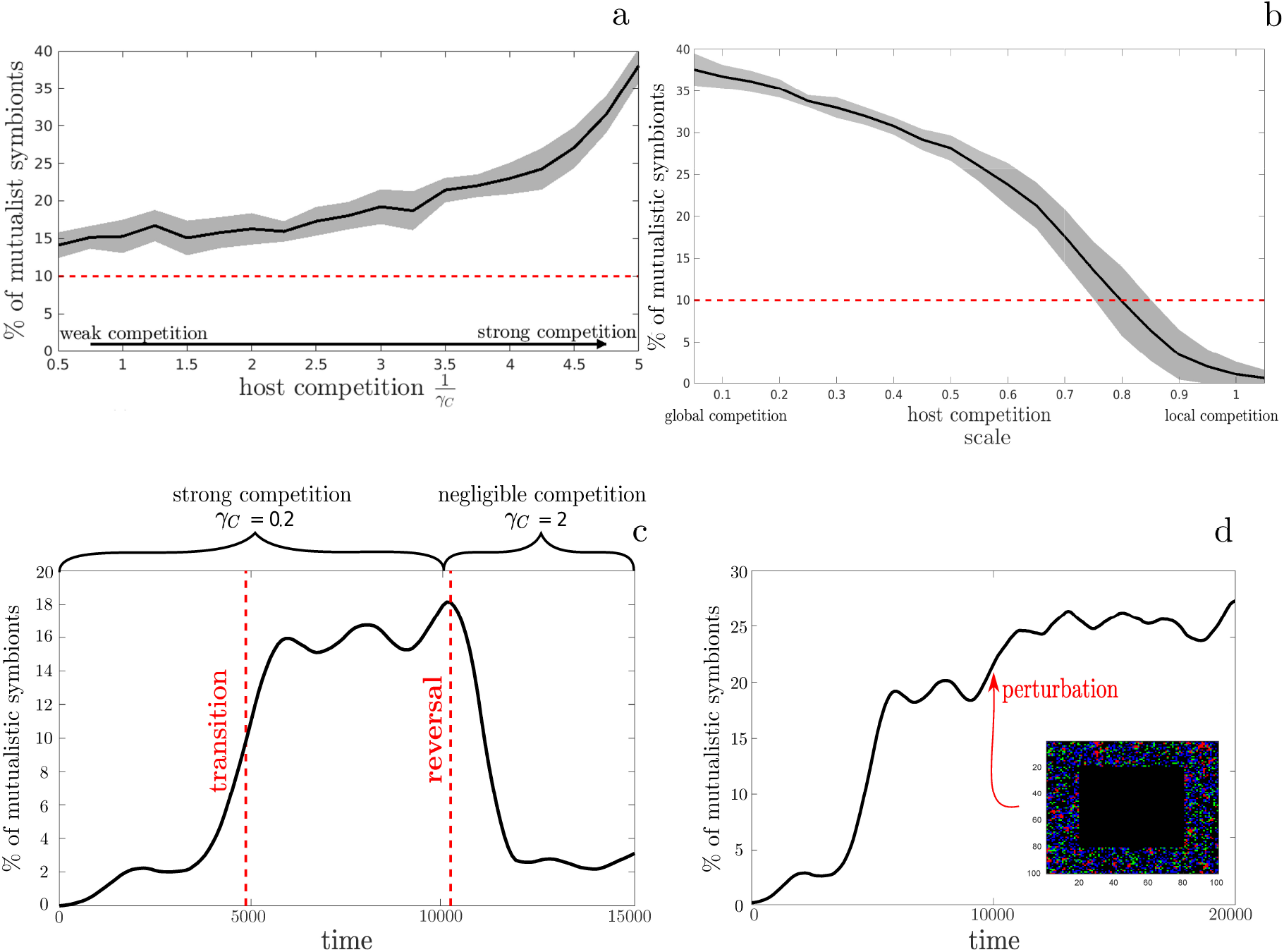
The role of host competition in the transition to mutualism. Effects of competition strength *γ*_*c*_ (panel a) and spatial scale *w*_*h*_ (panel b) on the percentage of mutualistic symbionts with dispersal cost *d* = 0.45 and after 10^4^ time steps and over 50 replicates. Black curves is the median and shaded regions are 95% confidence intervals. Dashed red line is the transition threshold of 10%. In panel a) competition is global (*w*_*h*_ = 1) and in panel b) competition is strong *γ*_*C*_ = 0.2. Panel c) represents the effect of competition on the transition and on the maintenance of mutualism (simulation without dispersal cost *d* = 0). Panel d) presents the effect of a reduction in competition caused by a perturbation eradicating all organisms in a large square. The perturbation occurs around *t* = 10^4^, which is 5000 time steps after the transition. Other parameters are *m* = 0.06, 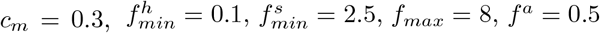 and *β*_*max*_ = 0.5.

Another way to investigate the effect of competition is to reduce host density, through the eradication of hosts in a region after a transition to mutualism. At first, the perturbed region is mainly recolonized by hosts and parasitic symbionts (Figure A10b), but mutualistic symbionts persist in the landscape. Due to the relaxation of global competition, the probability of host establishment is better, and the mutualistic clusters outside the perturbation zone gain in size, which explains why the proportion of mutualistic symbionts increases slightly despite the recolonization of the centre by parasites (Figure A10a). In the end, once recolonization is complete, the system returns to an equilibrium state whose trait distributions are close to distributions before the perturbation (details not shown). A similar experiment with a perturbation causing the death of 50% of uniformly occupied cells leads to the same results.

### Host dependency

Under favourable conditions leading to the transition to mutualism, the population of mutualistic hosts always persists in the absence of mutualistic symbionts, which excludes any absolute dependency of hosts for symbionts. However, *ecological dependency* may occur, where isolated hosts may have a negative growth rate because of intraspecific competition, although they would be able to form stable populations at lower densities. The transition to mutualism co-occurs with an increase in host density and thus an increase in intraspecific competition. If this increase in competition is sustainable only in the presence of mutualistic symbionts, the hosts are *ecologically dependent* on the symbionts. In order to determine the occurrence of ecological dependency, the intensity of intraspecific competition between hosts was measured in a system at equilibrium after the transition to mutualism (e.g., at the end of Figure 2c), and subsequently used as a fixed parameter to test if the host population can now survive in the absence of mutualistic symbionts and mutation. We found that ecological dependency occurs in the hatched area of Figure 6, when the system evolves toward a mutualistic system in which the percentage of mutualistic symbionts is sufficiently large.

**Figure 6:**
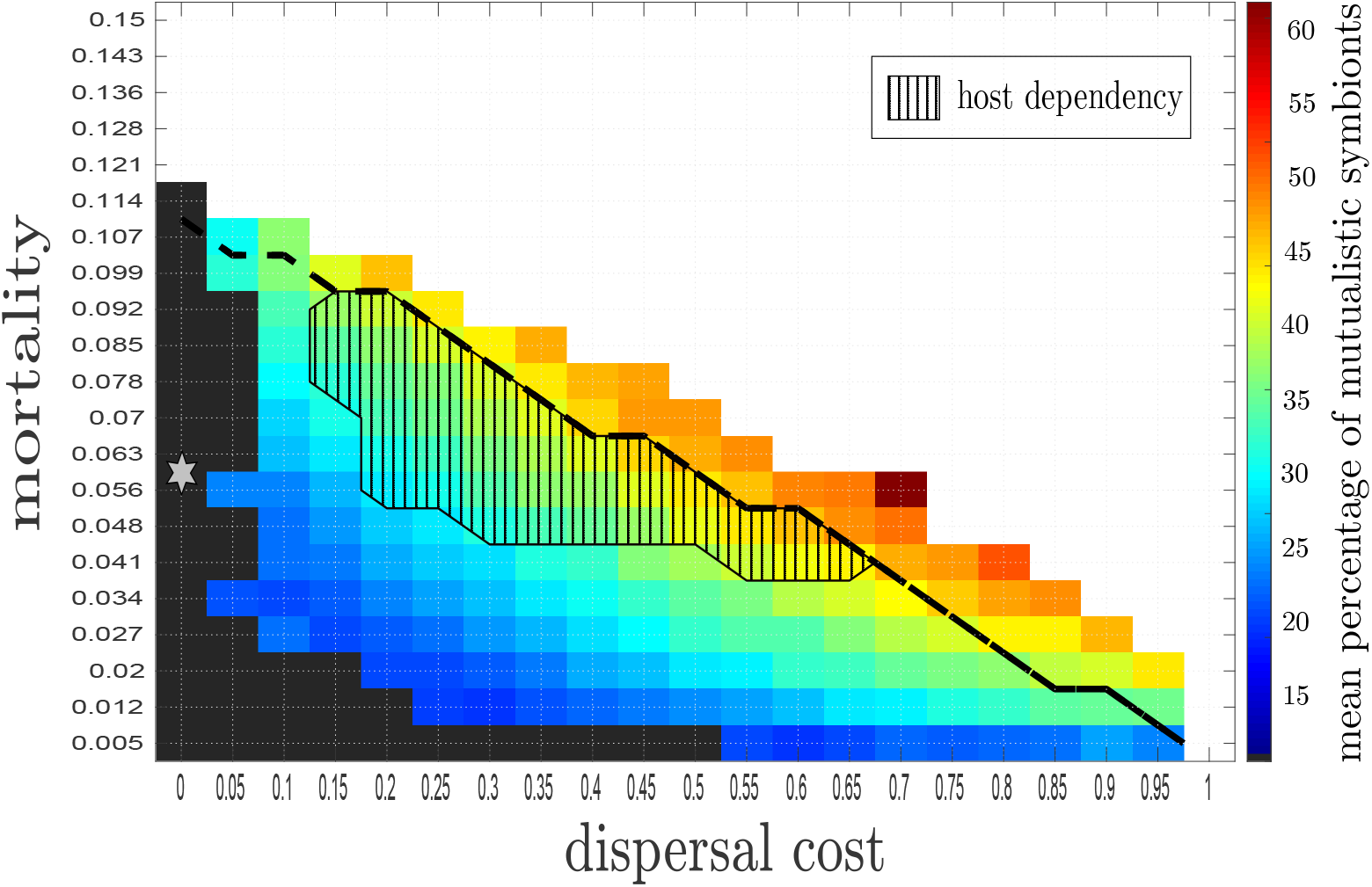
Percentage of mutualistic symbionts as a function of intrinsic mortality *m* and dispersal cost *d*. We run 50 simulations per parameter combination, with strong global competition *γ*_*C*_ = 0.2 and *w*_*h*_ = 1 with a maximum projection time of 10^4^ steps. The percentages are averaged over the simulations leading to transition, if any occur. Above the black dotted line, the parasitic system is not viable, although the evolution of mutualism can occur above this line (evolutionary rescue). White cells correspond to the nonviability domain for the whole system, even with evolution. In the dark grey area, none of the simulations gave birth to transitions. The evolution of host ecological dependency occurs in the hatched area, where an average isolated host has a negative growth rate because of intraspecific competition. The grey star corresponds to the restrictive conditions of Figure 2. Other parameters are 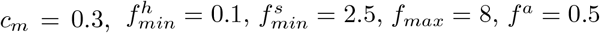 and *β*_*max*_ = 0.5.

Figure 6 further shows that both dispersal cost and mortality promote mutualism. For the parameter pair in the area indicated by the grey star, where dispersal cost is zero, Figure 2a showed that the probability of transition during the 10^5^ time steps is only 0.086, with a mean transition time of 7.10^4^. This explains why no transition occurred in Figure 6, where 50 simulations per parameter combination were performed, with only 10^4^ time steps. Finally, Figure 6 also shows that for some parameter combination, mutualism evolves even though the parasitic system is initially unviable. The viability of the parasitic system was assessed by simulations of 5000 time steps, without evolution. This implies that in a relatively short period of time in comparison to the transition times shown in Figure 2 for other parameter values, transitions can occur quickly enough and prevent the extinction of a parasitic system otherwise unviable. However this occurs rarely; Figure A13 shows that for some parameter combinations up to 90% of the simulations go extinct, the remaining being able to persist thanks to the evolution of mutualism. In those cases the mean percentage of mutualistic symbionts is much higher, ranging from 35 to 60%.

## Discussion

### The mechanisms underlying the transition to mutualism

In line with our main hypothesis, our results indicate that the transition from parasitism to mutualism occurs when mutualistic efforts evolve together with dispersal, despite the absence of vertical transmission or partner control. The following paragraphs review the mechanisms which contribute to the transition, and related them with the hypothesis formulated earlier.

#### The formation of clusters

Before the transition, a host performs better when alone; therefore, it has no interest in increasing its mutualistic effort and natural selection keeps it as low as possible. In contrast the symbiont population is limited by the number of available hosts, which increases when the symbiont becomes mutualistic. If mutualistic symbionts would help globally dispersing hosts, they would be counter-selected. However, in spatially structured populations rare mutants can interact with each other (Lion and Van Baalen, 2008), so if by chance mutations produce a mutualistic symbiont dispersing locally and interacting with a host dispersing locally as well, its offspring will benefit from the increased density of hosts in their neighbourhood and will form a mutualistic cluster (in line with hypothesis H1, Figures 4c) and A7). The cluster can then be invaded by parasitic symbionts dispersing globally, resulting in a dynamic equilibrium between mutualism and parasitism (in line with hypothesis H2, Figure 2c). Parasitic symbionts become themselves aggregated (Figure A7) since they develop around the mutualistic clusters, at their expense (Figure 4c). Joint evolution between mutualistic effort and dispersal results in a negative correlation between mutualism intensity and global dispersal (80% of mutualists disperses locally, Figure 3), which mirrors the link between altruistic behaviour and local dispersal (Koella, 2000; Le Galliard et al., 2005; Hochberg et al., 2008; Purcell et al., 2012; Mullon et al., 2018; Eldakar et al., 2010) as well as the relationship between local interactions and avirulence evolution (Boots and Mealor, 2007).

#### The key role of intraspecific competition

The invasion of a mutualistic cluster by parasites may cause its extinction and hinder the transition. We postulated that the higher fecundity of mutualistic clusters could compensate for their susceptibility to parasites (hypothesis H3). We instead found that, in the absence of dispersal cost, an eco-evolutionary feedback involving intraspecific competition between hosts was necessary for the transition. Indeed when competition between hosts is weak, the transition to mutualism rarely occurs

(Figure 2a), and when it does, the percentage of mutualistic symbionts remains low (Figure 5a). Conversely, when hosts strongly compete for resources, the ecological conditions change dramatically. The formation of mutualistic clusters (Figure 4a) increases population densities (Figures 2b and 2c), which enhances competition between hosts. Areas dominated by hosts associated with parasitic symbionts were initially viable, but their population growth rate becomes negative following the increase in competition. This creates empty space that can be colonized by mutualists, which still disperse globally from time to time. By lowering the abundance of parasitic symbionts, this also reduces the frequency at which mutualistic clusters are invaded by parasites. However, the key role of intraspecific competition only occurs when it is partly global (Figure 5b); if purely local mutualistic clusters cannot influence the viability of parasitic regions and will suffer from kin competition. In line with hypothesis H4, local competition between hosts for resources thereby prevents the transition to mutualism. Local competition between hosts for available space also occurs when hosts disperse locally, but this does not jeopardize the transition. Several obstacles must be overcome (simultaneity of the mutations, demographic stochasticity, possible invasions by parasites) before the mutualists are numerous enough to induce the shift in host competition, which explains why the transition needs some time to occur (Figure 2a). In sum, contrary to hypothesis H3 the transition is not directly caused by the higher fecundity of mutualistic pairs (which would fit soft selection, Wallace, 1975) but only indirectly by the increase in host competition, which renders areas dominated by parasites unviable (hard selection).

Empirical work has shown that the outcome of interactions between hosts and symbionts depends not only on the traits of the protagonists, but also on the surrounding ecological conditions (Bronstein, 1994). For instance, plants take advantage of seed-eating pollinators in the absence of alternative pollinators but not in their presence (Thompson and Cunningham, 2002). Mycorrhizae are beneficial for plants when soil resources are scarce while they are detrimental when resources are abundant (Johnson et al., 1997). In the above cases, the outcome of the interaction depends on both biotic and abiotic factors that are external to the host-symbiont system. Our model showed that the association with symbionts remains parasitic when host competition is low, while it evolves towards mutualism when host competition increases. In that case, the outcome of the interaction depends on intrinsic features of the interactions that are constructed by the eco-evolutionary dynamics of the system, as the emergence of mutualists increases host density.

#### The impact of dispersal cost and mortality

As expected, dispersal cost speeds up the transition (Figure 2a and 6) because it induces a selection pressure at the organismic level in favour with local dispersal, which increases the likelihood of the formation of mutualistic clusters. Mortality also enhances the probability of transition (Figure 6), but with another mechanism. We have stressed that competition between hosts creates an eco-evolutionary feedback loop, where the evolution of mutualism increases global densities, which strengthens competition and therefore turns the growth rate of the parasitic system negative. Given that mortality pushes the parasitic system towards its viability boundary, high mortality enhances the ability of competition to launch the transition. Although the transition occurs in a wide range of parameters where the parasitic system is viable, it is more likely when the parasitic system is close to extinction (Figure 6). However, mortality cannot itself trigger the transition since the parasitic system is unviable from the start when mortality is too high. Finally, mortality may also facilitate the transition through the reduction of global densities, which decreases the threat of parasites invading mutualistic clusters. The facilitation of mutualistic symbiosis in harsh environmental conditions has also been observed in previous empirical (Callaway et al., 2002; Maestre et al., 2003; Werner et al., 2015) and theoretical (Travis et al., 2006) works. However in the context of altruism the opposite relationship was found (Taylor and Irwin, 2000).

#### Evolutionary rescue

As evidenced by Figure 6, the evolution of mutualism can prevent the extinction of the parasitic system for parameter combinations that are just above the upper limit of its viability domain. This echoes the concept of evolutionary rescue (Ferriere and Legendre, 2013; Gomulkiewicz and Holt, 1995), according to which the persistence time of a population is longer with than without evolution. In the present case, instead of a single population, the populations of two distinct species are rescued by evolution. More generally, the parasitic system benefits from the evolution of mutualism even when it is initially viable, through an increase in population densities (Figure 2).

#### The evolution of mutualistic hosts

So far, only the mechanisms responsible for the evolution of mutualistic symbionts have been elucidated, but not those involved in the evolution of mutualistic hosts. Surprisingly, mutualistic hosts evolve after the transition (Figure 2c). Following the transition, the density of mutualistic symbionts is much higher, so that mutualistic hosts tend to be associated with mutualistic symbionts (Figure 4c), which disperse locally (Figure 3a). In that case, mutualistic hosts will increase the local density of mutualistic symbionts in the following generations, which will benefit their offspring provided that they disperse locally as well (Figure 3b). Symbionts may become less abundant for instance because of additional intraspecific competition between them, as in Appendix A.5. As a result, more hosts remain non-mutualistic because they are less often associated with a symbiont (Figure A12), which further highlights that the evolution of mutualistic hosts relies on high symbiont densities.

#### The role of quasi-vertical transmission

Although mutualistic symbionts are environmentally acquired, when both hosts and symbionts disperse locally this produces a similar effect as vertical transmission (as for mycorrhizae, Wilkinson, 1997), which we term “quasi-vertical” transmission. However, local dispersal (even 100%) is not equivalent to vertical transmission because host and symbiont offspring can disperse to any of the 8 neighbouring cells, so vertical transmission due to specific reproductive and physiological adaptations would have produced transitions to mutualism more easily. Moreover, the colonization of empty space by a mutualistic pair requires that both species disperse to the same remote place by chance, whereas in the case of vertical transmission this always occurs. Nevertheless, since hosts need to colonize empty space a significant fraction of hosts with mutualistic phenotypes also dispersed globally (∼ 40%, Figure 3), which partly counteracts the necessity of quasi-vertical transmission. As well as hosts, mutualistic symbionts may also suffer from limited dispersal when they need to percolate in a landscape of non-adjacent hosts, which explains why they maintain ∼ 20% of global dispersal (Figure 3). On the other hand, parasitic symbionts also evolve towards an intermediate dispersal strategy, although they tend to disperse globally much more often (∼ 80%, Figure 3). In purely parasitic systems it has been shown that some degree of vertical transmission, which is close to local dispersal in our case, is necessary for persistence in fragmented landscapes (Su et al., 2019; Schinazi, 2000). In those cases as well as here, the parasitic population needs some degree of local dispersal in order to exploit a patch of hosts, once it has been “found” by global dispersers. Intermediate dispersal strategies were shown to favor persistence of a variety of systems. For instance, frequent short-distance and rare long-distance dispersal together favor metacommunity persistence in fragmented habitats (Huth et al., 2015) and intermediate migration rate is required for the spread of cooperative strategies in spatial prisoner’s dilemma games (Vainstein et al., 2007).

The retention of some degree of global dispersal in both hosts and symbionts in order to colonize remote suitable places has another advantage; it indeed tempers local overpopulation generated by mutualism. Since overpopulation due to local dispersal increases kin competition, this reminds the evolution of altruism, which can be limited by kin competition (Wilson et al., 1992; Alizon and Taylor, 2008), as well as the evolution of dispersal which is in part due to the reduction of kin competition (Hamilton and May, 1977; Poethke et al., 2007; Harada, 1999). A mixed strategy combining both dispersal modes takes advantage of kin selection and simultaneously maintains the opportunity to escape kin competition. Figure 5b shows that purely local competition between hosts prevents the transition to mutualism because kin competition overcomes kin selection. Similarly, the evolution of cooperation by group selection can be hindered if competition between groups is local (Akdeniz and Van Veelen, 2020). In nature, global competition between hosts may arise when plants compete for water present in the same groundwater (Lejeune et al., 1999; Rietkerk et al., 2002), while competition for light is more local. Thus, the evolution of mutualism may depend on the dominant form of competition for resources between hosts.

### Assumptions, limitations and generality of the model

Our results rely on several hypothesis which have, if violated, either positive (vertical transmission, plastic costs) or negative (antagonistic coevolution, sexual reproduction, superinfections) effects on the likelihood of the transition to mutualism.

#### No vertical transmission

We excluded the possibility of vertical transmission because it is a complex feature involving many traits, which more likely evolve some time after the transition once the mutualistic relationship is well established. For this reason an alternative mechanism is needed, and our results demonstrate that the coevolution of mutualistic effort and limited dispersal in both species can mimic vertical transmission, as argued above. However, in some parasitic systems (e.g. birds displacing parasitic flatworms, ticks carried on large vertebrates) vertical transmission may be a passive feature, present from the start. In such cases the evolution of mutualism is theoretically possible even if hosts keep dispersing globally, provided that mutations turning the parasites into mutualists exist.

#### Benefits and costs of mutualism

Benefits only depend on the interaction trait of the partner. In turn, costs depend on the interaction trait of the focal organism as well as on the benefits provided by the partner (Appendix A.1). This would correspond for instance to the development of organs like plant domatia (Szilágyi et al., 2009): if the symbiotic ants are mutualistic, the plant can grow bigger, thereby producing more domatia, which is more costly in absolute terms. An alternative would be to assume that the costs do not increase with the mutualistic benefit; this would in any case be favourable to the evolution of mutualism. Moreover, in the model some cost is paid even if the partner is parasitic or if the host is free of symbiont. For instance, domatia or extrafloral nectaries are unconditionally produced (Bronstein, 1998), even though domatia size can be plastic (Kokolo et al., 2020). Also, plants produce costly floral displays even in the absence of pollinators. Finally, another alternative arises when partners interact repeatedly, for instance during their growth. Using an iterated prisoners’ dilemma model, Doebeli and Knowlton (1998) assumed that large received benefits trigger higher investment in the relationship. The interaction traits therefore become subject to phenotypic plasticity, in function of the partner’s trait. This assumption favors the transition to mutualism since mutualists benefit more from being associated with mutualists. In contrast, our set of assumptions is more conservative.

#### Antagonistic coevolution of the parasitic system

The evolutionary dynamics of the parasitic system have been ignored here, although they might affect the probability of transition. In the model the hosts cannot become resistant against the parasitic symbiont, which fits with the “superpathogen” of the gene-for-gene model (Salathé et al., 2008). This can be interpreted as a monomorphic long-term result of Red Queen dynamics, some constrain preventing the appearance of new resistant and virulent alleles. However, if the host-parasite interaction is instead ruled by a matching allele model (Salathé et al., 2008), dispersal and the associated spatial structure is likely to maintain polymorphism (Sasaki et al., 2002). During the early stages of the transition, formerly parasitic symbionts turned mutualistic will inherit this matching genetic system and will need to find compatible hosts. This adds another requirement, rendering the transition less likely.

#### Asexual reproduction

Many models of (co)evolutionary dynamics assume asexual reproduction (e.g. Kéfi et al., 2008; Loeuille and Loreau, 2005), especially within the framework of Adaptive Dynamics (e.g. Dieckmann et al., 1995; Loeuille and Loreau, 2005). In the case of sexual reproduction, recombination may soften the correlation between dispersal and interaction traits, which is nevertheless essential to the transition. However, the work of Dieckmann and Doebeli (1999) on the coevolution between a niche and a mating trait showed that linkage disequilibrium can itself evolve, thereby preserving the correlation between traits. In the present case a linkage disequilibrium between dispersal and interaction traits may also evolve; we therefore speculate that sexual reproduction would not prevent the transition in the long term, but only delay it.

#### Superinfections

We previously assumed that only a single symbiont could infect a host, however several strains may compete within the same host (Bongrand and Ruby, 2019; Zytynska and Weisser, 2016; Alizon et al., 2013). The host may be able to prevent the proliferation of parasitic strains (Sachs et al., 2010b), but parasitic strain may also overcome the others, which could prevent the evolution of mutualism (Jones et al., 2012). An extension of the model, presented in Appendix A.7, includes superinfections where mutualistic symbionts can be dislodged by parasites reaching the same host. When superfinfection probability rises above 50%, the transition is prevented, otherwise mutualistic symbionts can persist, although at lower densities than without superinfections (Figure A14). Thus, although superinfections are clearly detrimental to the transition, mechanisms favouring the evolution of mutualism in our present model can resist some degree of competitive exclusion by parasites.

#### The evolution of cheating

Our main interest was to understand how mutualism can evolve from a parasitic relationship (Roughgarden, 1975; Drew et al., 2021) but mutualism may also have evolved in the first place, the classic evolutionary problem in this case being how can it resist to the invasion of “cheaters” (e.g. Sachs et al., 2010a; Jones et al., 2015; Ferriere et al., 2002). According to Jones et al. (2015), cheating “(1) increases the fitness of the actor above average fitness in the population and (2) decreases the fitness of the partner below average fitness in the partner population”. The latter condition is always satisfied by parasitic symbionts, but the former remains to be checked. Simulations starting with mutualistic symbionts only are rapidly invaded by parasites, leading to an evolutionary equilibrium identical to the one reached by Figure 2c (details not shown). The population-level fitness (sensu Metz et al., 1992) of parasites is therefore positive when they are rare, thereby satisfying condition (1), and it gradually decreases to zero until the evolutionary equilibrium is reached. Hence, our model also accounts for the invasion by cheaters of an initially mutualistic system, leading to a coexistence of both strategies. Mutualism may also evolve from a competitive interaction, if two competitors start exchanging resources, each being a better exploiter of the resource it provides, and limited by the resource it receives (De Mazancourt and Schwartz, 2010). However it is unknown to what extent this kind of mutualism is sensitive to cheating; spatial effects similar to those studied here might stabilize it.

### The interplay between several levels of selection

Although the first models of group selection relied on well-defined groups (e.g. Wilson, 1975; Smith, 1964), multilevel selection theory has since been extended to fuzzy group boundaries and more complex landscapes (e.g. Lion and Van Baalen, 2008; Nunney, 1985; Tekwa et al., 2015) like in the present case. Earlier in the discussion, intermediate dispersal has been interpreted as the result of a balance between two components of inclusive fitness, kin selection and kin competition, which have been recognized as particular cases of multilevel selection (Goodnight, 2005; Lion and Van Baalen, 2008; Queller, 1992; Sober and Wilson, 1999). Although our model is too complex for an analytical derivation of inclusive fitness, this should be possible in principle, as it has been done for simpler models of the evolution of altruism (Hamilton and Fox, 1975; Lion et al., 2011; Lehmann et al., 2007; Marshall, 2011; Wade, 1980). However, the levels-of-selection problem is more a question about the level at which there is a *causal* link between character and fitness (Okasha, 2006, 2016; Sober, 1984; Sober and Lewontin, 1982), rather than the level at which a mathematical formulation of fitness can be derived (“bookkeeping” in the words of S. J. Gould 2002, p. 619). Following Sober (1984), we will consider that selection at a given level of organization occurs if the different entities belonging to this level are variable with respect to some property causally involved in the survival or reproduction of the organisms forming the entities. Since Sober’s formulation has been originally framed in the context of group selection, we first discuss how the levels-of-selection problem for mutualism can be related to the group selection debate in the context of altruism. The mechanism by which parasitic symbionts and hosts can invade mutualistic clusters is a two-species version of the tragedy of the commons (Garrett, 1968; Feeny et al., 1990; Hardin, 1998). In the case of altruism, the tragedy of the commons can be bypassed by local dispersal which triggers the formation of cooperative clusters (Mitteldorf and Wilson, 2000; Le Galliard et al., 2003; Eldakar et al., 2010), as in the present case. The evolution of altruism results from the conflict between two levels of selection, the organismic-level favouring cheaters and the group-level favoring altruism (Van Baalen and Rand, 1998; Simon et al., 2013; Wilson and Sober, 1989). At a given time step, neighbouring altruistic organisms help each other, which favors their fecundity. Shortly after the local density of altruists increases, which is beneficial for their offspring’s fecundity as well. Since the transition to mutualism is egalitarian whereas the transition to altruism is fraternal, it is unclear if the evolution of mutualism involves the same levels of selection as for altruism. Sure enough, mutualism is also counter-selected at the organismic level, since mutualism is costly to both hosts and symbionts. However, differences between altruism and mutualism may arise at higher organization levels because at a given time step mutualists help their heterospecific partners but not their neighbouring conspecifics. In the present model the evolution of mutualism involves selection at the level of the host-symbiont pair, since at a given time step the reproduction of each of its organisms depends on the properties of the pair (the interaction traits *α*_*h*_ and *α*_*s*_). This resembles the tit-for-tat strategy where cooperators are selected at the pair level (Wilson, 2004; Sober and Wilson, 1999). The mutualistic host-symbiont holobiont therefore emerges as a new unit of selection (Roughgarden et al., 2018; Drew et al., 2021).

Considering several times steps in a row, another level of selection appears. Since mutualists also disperse locally (Figure 3), after some time a mutualistic pair may trigger the formation of a mutualistic cluster (Figure 4c). Neighbouring mutualistic pairs do not help each other directly at a given time step, but indirectly by increasing the likelihood that their offspring will encounter mutualistic partners in the subsequent time steps. Although only hosts and symbionts reproduce in the traditional sense of organismic reproduction, the association between mutualistic hosts and symbionts is also re-produced (Doolittle and Inkpen, 2018; Griesemer, 2001) via local dispersal and cluster formation. Selection at the cluster level therefore occurs, since clusters dominated by mutualistic pairs will favour the reproduction of organisms and the re-production of mutualistic pairs. The re-production of pairs constitutes a another mechanism of inheritance, different from the one occurring during organismic reproduction, and fits with the idea that major evolutionary transition involve the evolution of informational systems (Szathmáry, 2015; Szathmáry and Smith, 1995). It is therefore hard to match Hull’s (1980) categorization of replicators (here, hosts and symbionts) and interactors (here, pairs), since during the transition mutualistic pairs also acquire a replicative power via the evolution of local dispersal. This also emphasizes that Sober’s (1984) formulation of group selection needs to be generalized for the present context, since the properties of clusters favor not only the reproduction of organisms but also the transmission of higher-level properties. Mutualistic clusters are self-perpetuating systems (Lenton et al., 2021), some of their properties being homeostatic (Ibanez, 2020). However, we believe this is not enough to qualify to evolutionary individuality (sensu Godfrey-Smith et al., 2013) since conflicts are still vivid (Queller and Strassmann, 2016); mutualistic clusters being prone to the invasion by parasites (Figure 4c). The transition to mutualism may lead to genuine evolutionary individuals during later phases which would suppress conflicts.

Lastly, in the absence of dispersal cost mutualism rarely invades when host competition is weak (Figure 2a), despite the occasional formation of mutualistic pairs. Without dispersal cost, competition between hosts at the global scale is necessary for the transition to mutualism (Figure 5b). The global scale therefore constitutes another level of organization involved in the transition to mutualism. Global competition between hosts acts as an environmental factor mitigating selection at the different organization levels discussed above. This environmental factor is not fixed by a parameter but instead determined by the evolutionary dynamics of the whole system, it is at the same time subject and object of evolution (Lewontin, 1982, 1983).

### Host dependency and irreversibility of the transition

Major transitions in evolution are characterized by their irreversibility and by the interdependence between the agents (Szathmáry and Smith, 1995; Estrela et al., 2016). The model does not include any physiological or developmental dependence of the host on its symbiont, or any loss of functions in the host due to gene transfers, because we assumed that this generally occurs during later stages of the evolution of mutualism (Nguyen and Baalen, 2020). Instead, dependence has been defined from a population dynamics perspective: the host is *ecologically dependent* when its population growth rate is negative in the absence of the symbiont. In that case the host can produce offspring, although not enough to compensate for mortality. In line with hypothesis H5, we found that mutualistic hosts deprived of their symbiont exhibit a negative growth rate when the host density after the transition to mutualism becomes sufficiently large (Figure 6). This ecological dependency resulted from the density-dependent competition between hosts and the assumption that mutualism is costly for the host, even when its symbiont is absent (as discussed above). However, ecological dependency is not absolute: once the density of hosts becomes sufficiently low, the mutualistic hosts alone are viable. Dependency may become absolute for a sufficiently high cost of mutualism, but in these conditions the transition to mutualism will not occur.

If host competition strength decreases permanently, for instance following the continuous supply of extra resources, the reverse transition back to parasitism occurs (Figure 5c). This has been documented in nature as well (Pellmyr and Leebens-Mack, 2000; Kawakita et al., 2015; Sachs et al., 2011), although the mechanisms involved may well be different. Reversal towards parasitism occurs because ecological dependency relies on host competition, which change with host densities, highlighting that mutualistic symbiosis may be sensitive to environmental change (Drew et al., 2021). However, if host competition decreases punctually, e.g., following a perturbation of a fraction of the landscape, then mutualism persists (Figure A10) because mutualistic clusters take advantage of the reduction of global host competition to colonize free cells around them. This leads to an increase in host competition; in that case mutualism can restore the ecological conditions allowing its own persistence, as in a niche construction process (Lewontin, 1982, 1983; Odling-Smee et al., 2013; Laland et al., 2016). Niche construction is generally understood as the improvement of abiotic conditions (e.g. Arnoldi et al., 2020). In the context of mutualism, it is due to the improvement of host densities, which induces an increase in host competition. This also occurs at the beginning of the transition, when the first mutualistic clusters trigger an increase in global host density. Although this has not been tested formally, the reversion is also very likely to occur if host competition for resources shifts from global to local, since it is apparent from Figure 5b that local competition completely prevents mutualism, even in the presence of dispersal cost.

## Conclusion

In the present paper, we aim to understand the mechanisms promoting the transition from parasitism to mutualism. To tackle this issue, we develop an agent based model on a lattice. In our general model, we only assume that the mutualistic interactions influence the fecundity of both partners and that hosts face density-dependent competition; and ensure that the antagonistic system is stable in absence of mutations. We found that in the absence of vertical transmission or partner control mechanisms, the joint evolution between mutualistic effort and local dispersal can trigger the transition from parasitism to mutualism, provided that intraspecific competition between host is sufficiently global and that either dispersal cost or competition strength is large enough. Unexpectedly, we found that mutualistic clusters invade the antagonistic system thanks to their ability to increase the population densities of both partners, thereby triggering global competition between hosts and rendering regions where hosts are mainly associated with parasitic symbionts unsuitable. In contrast, the higher fecundity of mutualists is not advantageous enough to compensate for the ability of parasites to invade mutualistic clusters, contrary to our initial expectation. Our results suggest that the eco-evolutionary feedback involving competition between hosts might promote the transition from parasitism to mutualism in a wide range of biological systems, such as plant-fungi, plant-ant and plant-seed-eating pollinator interactions.

## Acknowledgements

We are grateful to Jean-François Arnoldi, Cédric Gaucherel, Sonia Kéfi, Eva Kisdi, Sébastien Lion, Nicolas Loeuille, François Massol (all listed by alphabetic order) and three anonymous reviewers for their thoughtful comments and discussion on this work.

## Data and code accessibility

All the codes used to compute the outcomes of our model and the figures of the paper are available on the following doi.org/10.5281/zenodo.6463210

## A Appendix

### A.1 Mathematical and numerical details of the model

We present here the mathematical underpinnings of the model as well as some details of the numerical computation.

#### Rules of the individual based model description

Our model follows the cycle presented in Figure A1:

- Host and symbiont die with fixed probability *m* ∈ (0, 1).
- They produce offspring, possibly with different traits from them due to mutation. The fecundity of the parents depends on their two traits (*α, ε*) ∈ [0, 1]^2^ and on their interactions with their possibly cell-sharing partner.
- The offspring are dispersed according to the parents’ dispersal traits *ε*.
- The offspring of the hosts may establish only in empty cells, while the offspring of the symbionts can only establish in cells already occupied by a solitary host. If several organisms arrive in the same cell, a lottery determines which one will establish, while the others die.

In our numerical computations, mutations occurred only after the descendant was successfully established in a cell. This procedure saves computational time and did not influence our results because offspring dispersal and establishment do not depend on their traits but only on their parent traits. Furthermore, the mortality process was applied to both types of agents simultaneously, while the reproduction and dispersal processes were applied consecutively to the hosts and then to the symbionts. We confirmed that the order of the algorithm did not qualitatively affect our results.

#### Fecundity and the average offspring number

The fecundity *f* of an agent depends on its mutualistic interaction trait *α* as well as the interaction trait of its cell-sharing partner. This continuous trait ranging between 0 and 1 determines the intensity of the agent investment in the mutualistic relationship.

We assumed a positive interaction trait dependence between agents. A mutualistic agent tends to increase the fecundity of its cell-sharing partner. The interaction fecundity 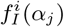 of an organism of type *i* ∈ {*h, s*}, (h = host, s = symbiont) interacting with an organism of type *j* ∈ {*s, h*} with trait *α*_*j*_ was defined by

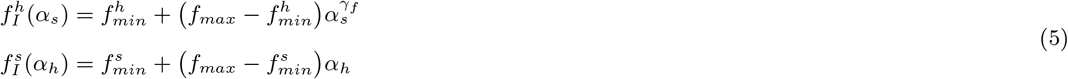

The coefficient *γ*_*f*_ corresponds to the mutualistic strength on the host fecundity. Using a coefficient *γ*_*f*_ *>* 1, we create a convex function allowing a transition from parasitism to mutualism for a central value of the symbiont interaction trait. However, note that modifying the shape of this fecundity curve (from concave to convex via linear) does not qualitatively change our results.

**Figure A1:**
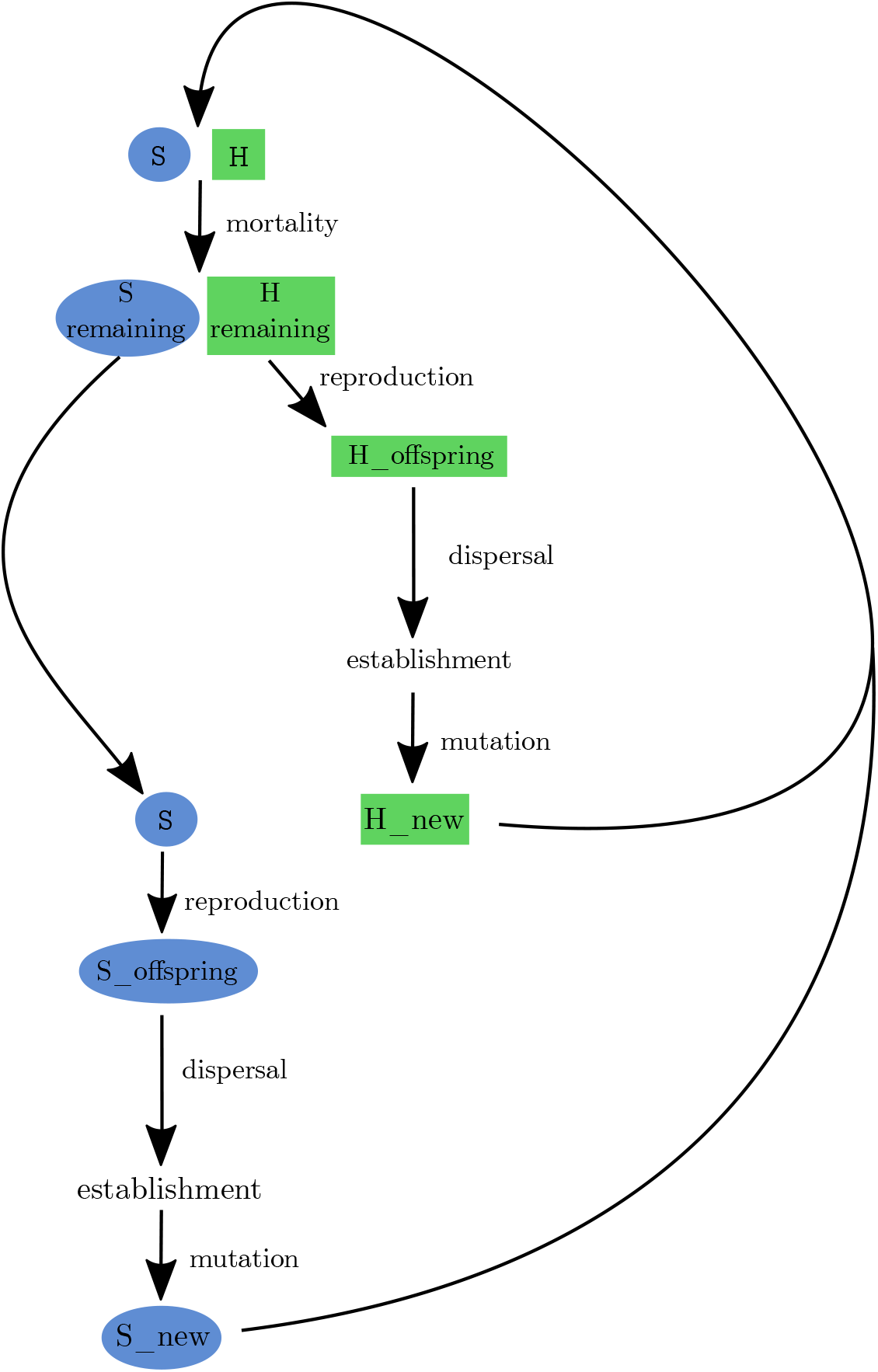
Sketch representation of the individual based model. The host population (H) and the symbiont population (S) undergo intrinsic mortality, then reproduction, dispersal, establishment, and finally mutation. The mortality step is simultaneous for the host and the symbiont, while the other steps occur first for the host and then for the symbiont.

On the other hand, a mutualistic agent has an intrinsic cost reducing its fecundity. The mutualism cost *C*_*m*_(*α*_*i*_) of an organism of type *i* ∈ {*h, s}* (h = host, s = symbiont) ranges between 0 and 1, and it increases with interaction trait *α*_*i*_ of the agent. It is defined by

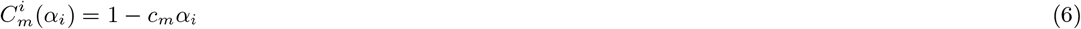

where *c*_*m*_ is the maximal cost of mutualism.

Thus, for the host as for the symbiont, the fecundity *f*_*i*_ of an organism *i* interacting with an organism *j* is the product of the interaction fecundity *f*_*I*_ (*α*_*j*_) defined by (5) and the cost of mutualism *C*_*m*_(*α*_*i*_) defined by (6).

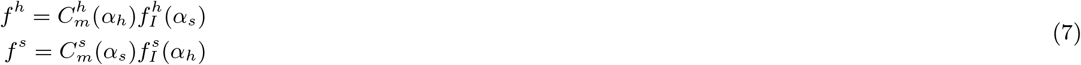

When a host agent is alone in a cell, its fecundity is defined by its intrinsic host fecundity *f*^*a*^ weighted by its mutualism cost *C*_*m*_(*α*_*h*_): Fecundity of the solitary host:

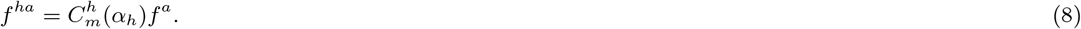

We assume that the cost of mutualism is paid regardless of whether the interaction is realized.

In general, the average offspring number is not integer, yet the number of offspring in our model can only be represented by an integer. Thus, in the numerical algorithm, the fecundity was used as the *λ* parameter of a Poisson distribution. If the value drawn from the distribution was greater than the maximum fecundity *f*_*max*_, then it was set back to the maximum fecundity.

#### Mutualism/parasitism threshold

In our model, the presence of a host always produces a net benefit for the symbiont. However, the presence of the symbiont might be detrimental for the host. Indeed, the fecundity of a host *h* interacting with a symbiont *s* is 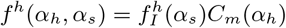, while the fecundity of the same host *h* without a symbiont is *f*^*ha*^(*α*_*h*_) = *f*^*a*^*C*_*m*_(*α*_*h*_). Thus, the host has net benefit only if its fecundity in association with a symbiont is larger than its fecundity alone. Therefore, mutualism only occurs when 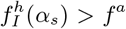. Otherwise, the interaction is parasitic. This criterion does not depend on the host mutualism trait *α*_*h*_ because hosts always pay the same mutualism cost. Thus, we can define the mutualism/parasitism threshold 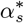 such that 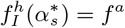; thus, we obtain

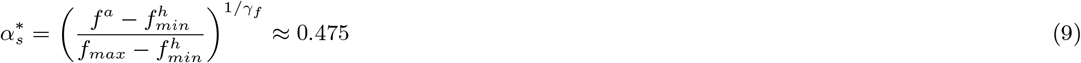

with the parameters set in Table 1.

#### Competition

To test the effect of the spatial scale of the competition, we introduced a scale parameter *w*_*h*_ ∈ [0, 1] that weighs the effect of local 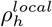 and global 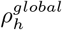 host density on the competition. The establishment probability thus satisfies

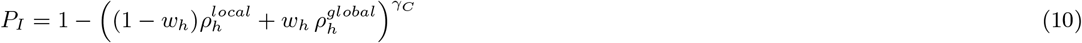

The local host density 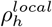 corresponds to the host density in the 8 neighbouring cells surrounding the implantation cell of the host, while the global density 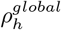 corresponds to the host density all over the landscape (see Figure A2 for a schematic representation).

**Figure A2:**
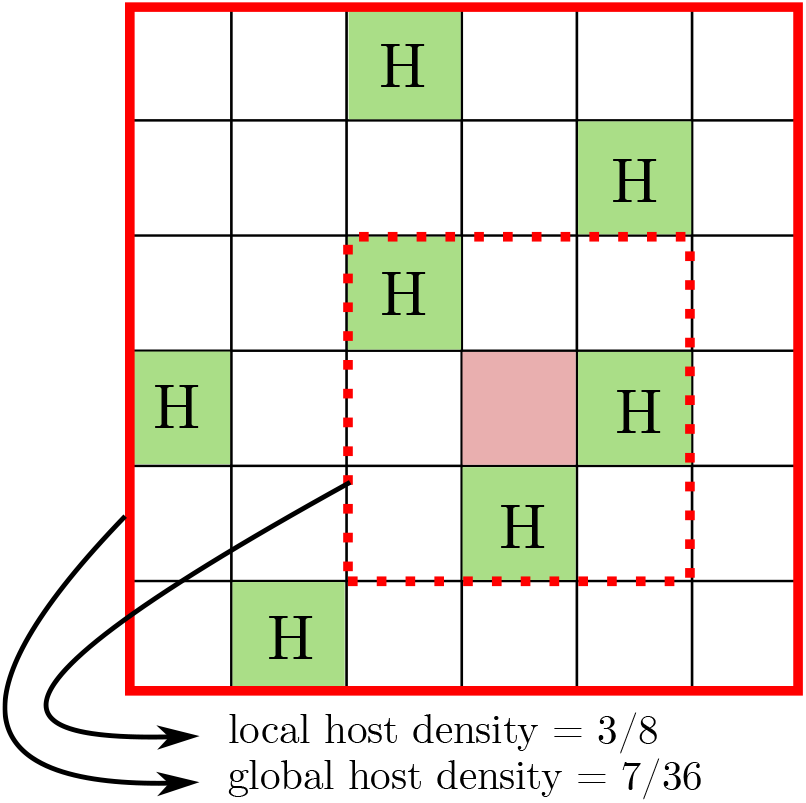
Local and global host densities influence the probability of establishment in the focus cell (pink filled square). The global density corresponds to the host density in the whole 36-cell landscape. The local density corresponds to the density in the eight cells vicinity around the focus cell.

These competition scales may have various ecological explanations. For instance, plants sharing the same water table face global competition for this resource. Conversely, the competition for light between plants is an example of local competition. Thus, our competition scale model allows us to describe the competition for several different resources that may appear at different scales. Following our previous examples, if the water supply represents 90% of the competition and light supply represents only 10%, then the competition scale *w*_*h*_ is *w*_*h*_ = 0.9 (90% global competition and 10% local competition).

#### Distribution of mutation effects

During reproduction, organisms generate offspring with traits that can deviate from their traits due to mutation. The effects of mutation on each trait are independent. However, the mutation effect does depends on the trait of the parent. For instance, an organism with trait *α* will give birth to an organism of trait *α* + *β* where *β* is drawn from a distribution with probability distribution function given by *K*(*β* | *α*), which depends on the trait of the parent *α* (Figure A3). In our model, we use a modified Beta distribution with shape parameters (1, 3) to describe the effects of the mutation. More precisely, for a parent of trait *α*, the effect of mutation is a random variable *β* defined by

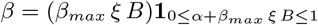

where *B* is a random variable, which follows a Beta distribution, *ξ* is a random variable independednt of *B*, which follows a Bernouilli distribution (**P**(*ξ* = 1) = **P**(*ξ* = −1) = 1*/*2). In other word, the random variable *β* follows the probability distribution function *K*(*β* |*α*), with *α* ∈ [0, 1]:

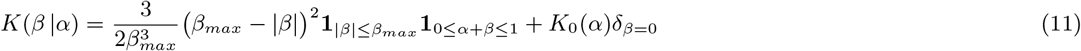

where **1** is the indicators function, *δ* is the Dirac mass and the function *K*_0_(*α*) is defined by

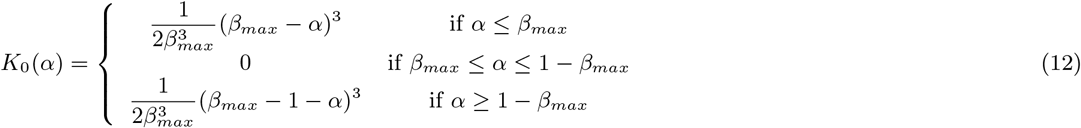

**Figure A3:**
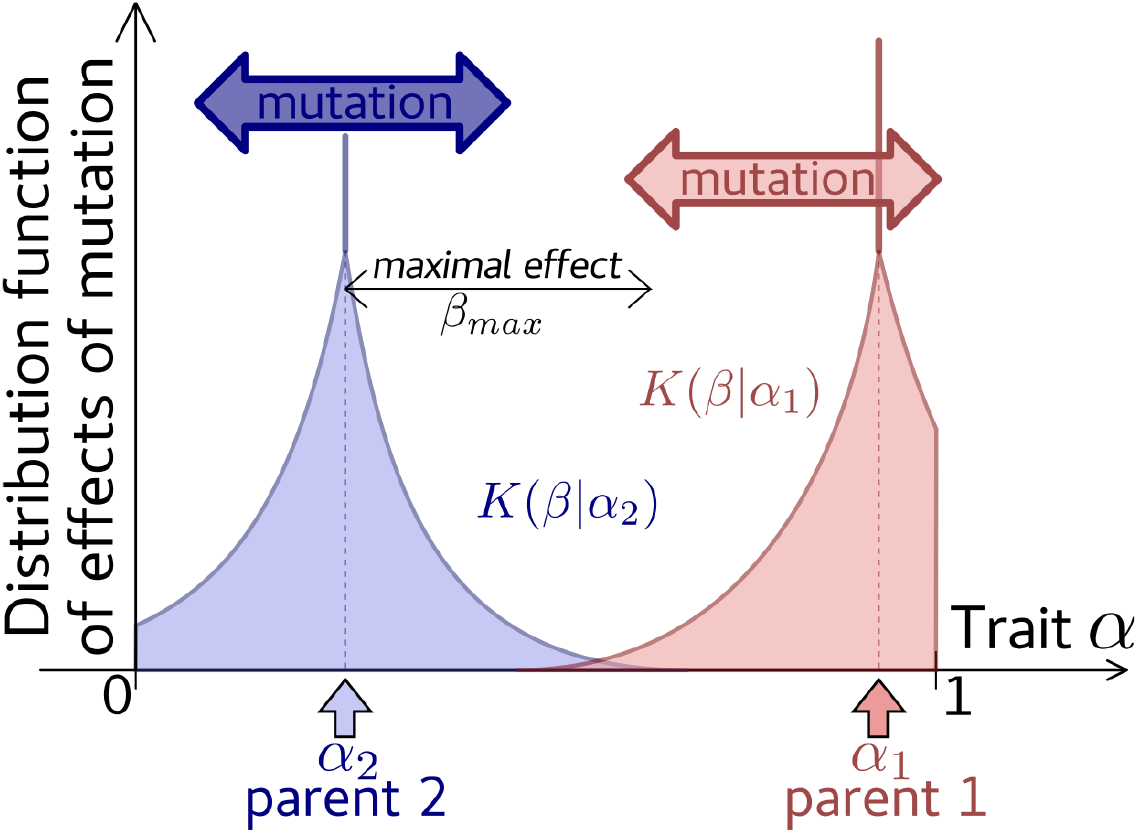
Distribution of mutational effects *K*(*β* | *α*). Each parent of trait, e.g. *α*_1_ or *α*_2_, produce offspring with trait *α*_*i*_ + *β* where *β* has the density *K*(*β* | *α*_*i*_) depending on its parent traits (red and blue curves for *α*_1_ and *α*_2_, respectively).

Moreover, we investigate the effect of the maximal effects of mutation *β*_*max*_ on the proportion of mutualistic symbionts. From our formula, we know that the mean effect of mutation depends on the trait of the parent *α* but it is proportional to *β*_*max*_, and it ranges between 3*β*_*max*_*/*8 for parents with intermediate trait (*α* ∼ 0.5) and 3*β*_*max*_*/*4 for parents with trait either close to 1 or 0. We show in Figure A4 that increasing the mean effect of mutation increases the proportion of mutualistic symbionts in the population. Thus large effects of mutation favour the emergence of mutualism. In our simulations we fix the maximal effect of mutation to *β*_*max*_ = 0.5.

#### Dispersal

At each time step, hosts and symbionts produce offspring which can disperse over the landscape either locally or globally. For each agent, the proportion of its offspring dispersing globally is given by the dispersal trait *ε*. The location of offspring dispersed locally is chosen randomly uniformly over the 8 neighbors of its parents, while the location of those dispersed globally is chosen uniformly over the entire landscape expected the location of the parent (Figure A5 for the description of the local and global scale). In particular, a globally dispersed organism can arrive in the local neighbor of parents as the locally dispersed one. Moreover, the offspring are dispersed independently from each other and their location is chosen independently of the current landscape. In particular, offspring can arrive at an already occupied location and symbionts’ offspring are not only dispersed in location where there is already an host. For instance if a host disperse 2*/*3 of its offspring at large distance from it, its dispersal trait satisfies *ε* = 2*/*3. Then the 2*/*3 of its offspring are dispersed randomly uniformly in the entire landscape (red stars in Figure A5) while the remaining 1*/*3 is dispersed locally around it (red circles in Figure A5).

**Figure A4:**
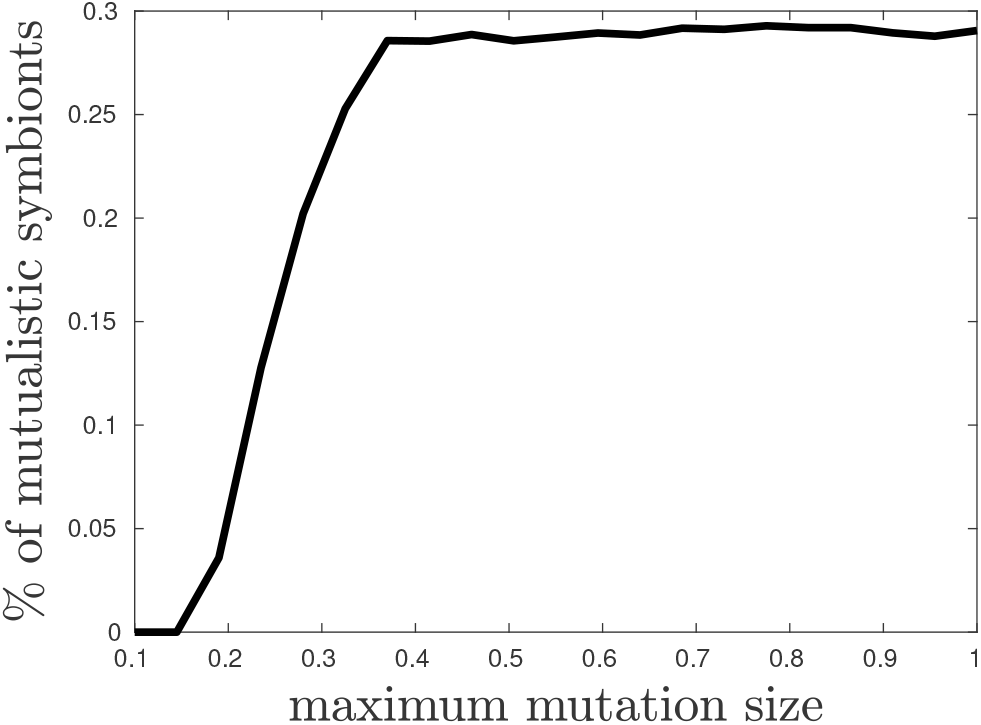
Effect of the average mutation effect (parameter *β*_*max*_ of distribution kernel of mutation effect) on the proportion of mutualistic symbionts.

**Figure A5:**
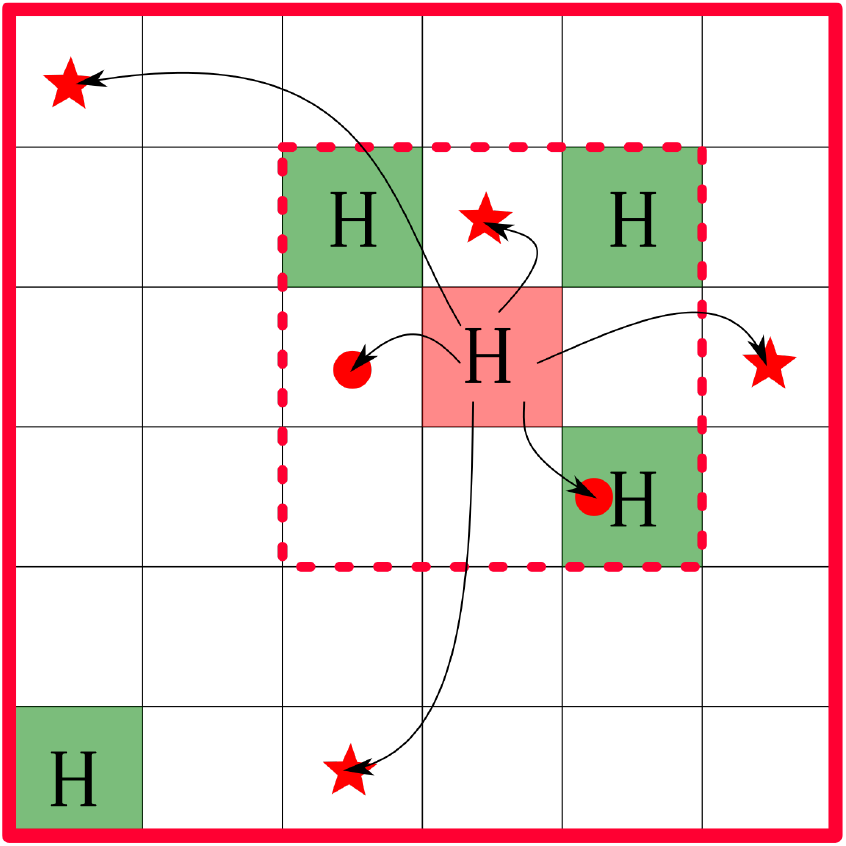
Local and global dispersal of offspring from the host in the red cell over the landscape. Local dispersal (red circles) occurs only within the neighborhood of the host (red dashed square) while global dispersal (red stars) occurs over the entire landscape (red plain square). The host has a dispersal trait *ε* = 2*/*3 and it disperses 6 offspring: 4 globally (red stars) and 2 locally (red circles).

#### Assortment index

To compute the assortment index, we measured the similarity between spatially neighbouring phenotypes for the spacial repartition resulting from the transition to mutualism and for the same spacial repartition but with phenotypes randomly redistributed among organisms. The assortment index corresponds to the difference between the measurement made on the space resulting from the transition to mutualism and the measurement on the randomly rearranged space. If the index shifts positively (resp. negatively) from zero, it means that similar phenotypes are closer (resp. more distant) than different phenotypes compared to random spatial distribution. This methodology is similar to that used in Pepper and Smuts (2002) and Pepper (2007).

##### Intraspecific assortment index

More precisely, for the intraspecific assortment index we use the following similarity index for host and symbiont. For each simulation and time *t*, we compute the similarity indices *S*_*h*_ and *S*_*s*_ respectively among hosts and symbionts, as follows

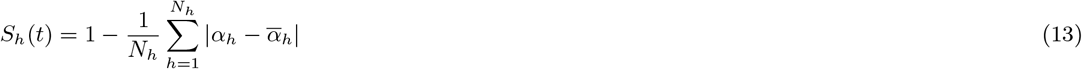

where *α*_*h*_ is the trait of the host *h* and *N*_*h*_ is the total number of host in the landscape at time *t*. The quantity 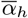 is the average trait in the neighborhood *V*_*h*_ of the host *h*. The neighborhood *V*_*h*_ of a host *h* is the 8 closest cells surrounding it (figure A2). It is defined by

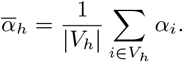

The similarity index among symbionts *S*_*s*_ is computed similarly.

Then for each time, we reshuffle the traits among the location occupied by hosts and symbionts and we compute the associated similarity indices using equation (13). We average those indices over 1000 replicates to compute the similarity indice *S*_*rh*_ and *S*_*rs*_ corresponding to a random spatial distribution.

Finally, We build the assortment index *A*_*h*_ as the difference between the similarity index of host *S*_*h*_ observed and the similarity index *S*_*rh*_ of host when we randomly assigned trait of the host over the landscape,

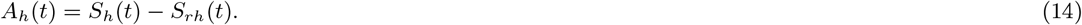

We also compare our assortment index with the spatial autocorrelation Moran index for the host and symbiont. The two indices show the same pattern. A positive spatial autocorrelation is observed after the transition occurred (Figure A6).

##### Interspecific assortment index

For the assortment index between host and symbiont, we also use a measure of similarity between the host and symbiont trait at each location of the couple. More precisely, we define for each simulation and each time *t* the similarity index *S*_*sh*_ between host and symbiont sharing the same location as follows

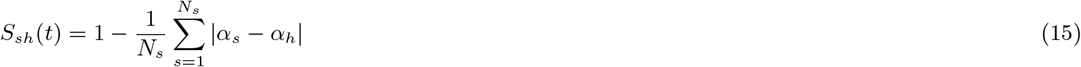

where *N*_*s*_ is the number of symbiont, which is also the number of host–symbiont couple. As before, we compare this observed index with the random index *S*_*rsh*_ defined by randomly rearranging pairs of symbiont and host and taking average over 1000 replicates. The assortment index *A*_*sh*_ is thus given by

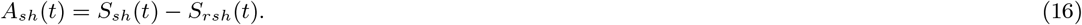

We also compare our index with the correlation coefficient between the interaction traits of hosts and symbionts. We find a positive correlation between trait in a same location (Figure A6).

**Figure A6:**
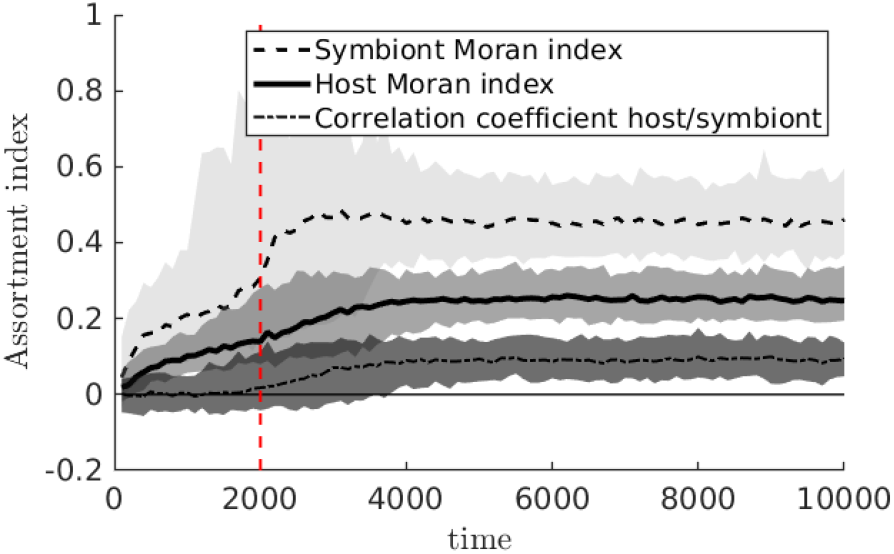
Spatial autocorrelation among hosts (plain curve) and symbionts (dashed curve) are described by the Moran index. The spatial correlation between the host and symbionts are described by the correlation coefficient (dash-dotted curve). The shadow regions corresponds to the 95% confidence interval and curves corresponds to the median over 100 replicates. The parameters are similar as Figure 4.

#### Aggregation index

From the assortment index analysis, we show that the symbionts and hosts are spatially assorted according to their trait. Now we aim to investigate how they are aggregated in space. We use a relative aggregation index based on a measure of the number of pair of neighbors. More precisely, we define for any spatial configuration the number of pairs of neighbors *P* where a neighbor of an organism is its 8 closest cells. For instance, Figure A2 provides a schematic representation of a host spatial configuration and the dashed square represents the neighborhood of the red organism. The number of pair of the red organism is 3 in this example. Then for any spatial configuration with *n* organisms, we can define the maximal number of possible pair of organism which is given by 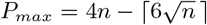 (Harary and Harborth, 1976). Thus, we define the aggregation index *𝒜* as the ratio between *P* and *P*_*max*_:

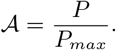

We compute the aggregation index over time for the hosts, the parasitic symbionts 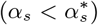 and the mutualistic symbionts 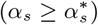 (Figure A7).

Hosts are always more aggregated than symbionts. Moreover, after the transition occurred, mutualistic and parasitic symbionts have the same spatial signature in terms of aggregation. This pattern was already observed in Figure 4 where we see mutualistic clusters surrounded by parasitic clusters.

**Figure A7:**
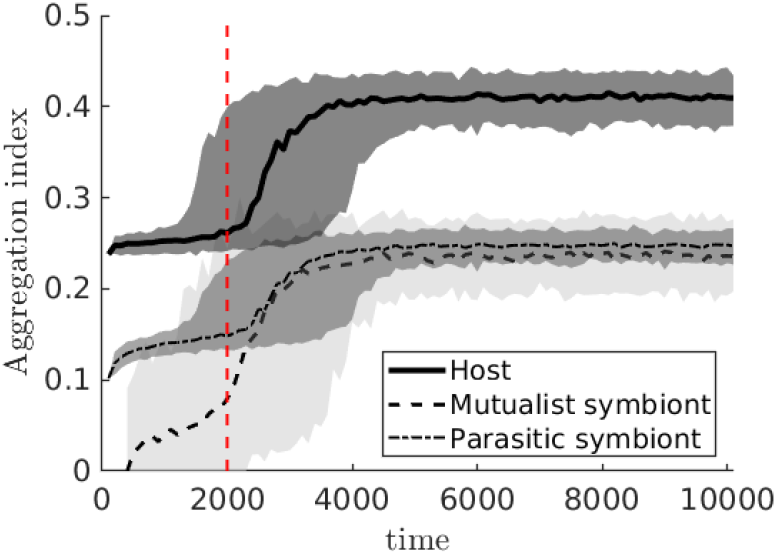
Aggregation index of the spatial distribution of hosts (plain curve), parasitic symbionts (dash-dotted curve) and mutualistic symbionts (dashed curve) over time. The shadow regions corresponds to the 95% confidence interval and curves corresponds to the median over 100 replicates. The parameters are similar as Figure 4.

### A.2 Mathematical approximation

In order to provide some heuristics about our stochastic model, we develop a simple deterministic model, with a monomorphic population of host and symbiont. The mathematical analysis also provides quantitative insights on our choice of parameters.

More precisely, we aim to describe the expected proportion of sites occupied by a monomorphic population of hosts and symbionts at equilibrium with interaction traits *α*_*h*_ and *α*_*s*_ respectively. We assume no mutations of interaction or dispersal traits and hosts and symbionts disperse globally randomly over the landscape composed of *N* sites. According to our stochastic model (see Fig. A1), the dynamics of the proportion of sites occupied by the host alone *ρ*_*ha*_ or host with symbionts *ρ*_*hs*_ is given by

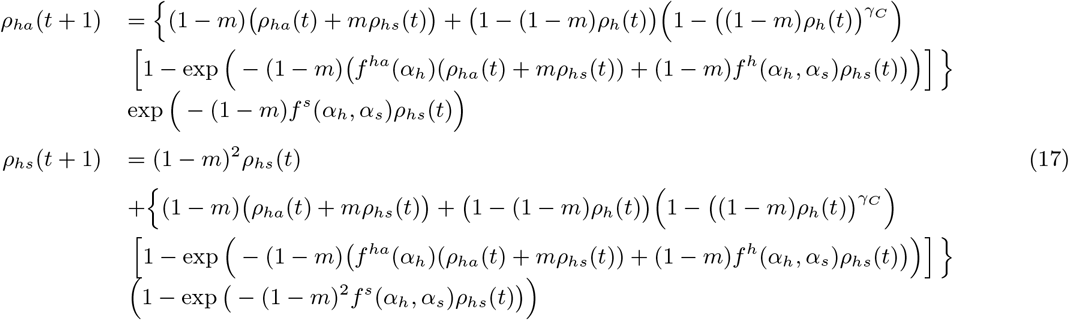

where *ρ*_*h*_ = *ρ*_*ha*_ + *ρ*_*hs*_ is the total proportion of hosts. Since hosts and symbionts first face mortality with rate *m*, the proportion of host alone becomes (1 − *m*)(*ρ*_*ha*_(*t*) + *mρ*_*hs*_(*t*)), where *mρ*_*hs*_ corresponds to hosts that have lost their symbiont, and the proportion of hosts with symbionts is (1 − *m*)^2^*ρ*_*hs*_(*t*). Then hosts produce offspring at a rate that depends on their partner: *f*^*ha*^(*α*_*h*_) (host alone) or *f*^*h*^(*α*_*h*_, *α*_*s*_) (host with symbiont). The total number of offspring is thus given by

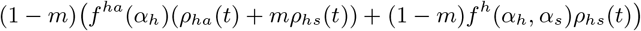

Since offspring are dispersed randomly uniformly over the landscape, the probability that at least one offspring enters a cell is given by

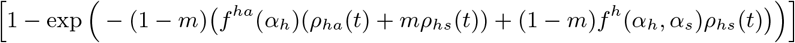

However, they can only colonise empty cells, whose proportion is (1 − (1 − *m*)*ρ*_*h*_). Moreover, once they enter an empty cell, their probability to establish in this cell depends on the host density *ρ*_*h*_ and it is given by 1 – ((1 − *m*)*ρ*_*h*_(*t*)) ^*γC*^). Finally, the symbionts produce offspring at a rate *f*^*s*^(*α*_*h*_, *α*_*s*_). Their offspring can only colonise alone host whose proportion is now given by the term between brackets. Since symbiont offspring are also randomly dispersed, the probability to invade a host alone is given by (1 − exp (− (1 − *m*)^2^*f*^*s*^(*α*_*h*_, *α*_*s*_)*ρ*_*hs*_(*t*))).

In this model, the traits are fixed – if 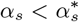 symbionts are parasitic while there are mutualistic if 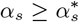. Since the symbionts need host to survive, the proportion of sites occupied by symbionts is *ρ*_*hs*_. Even if hosts and symbiont does not share the same mortality rate *m*, the model holds true by replacing *mρ*_*hs*_ and (1 − *m*)*ρ*_*hs*_ by *m*_*s*_*ρ*_*hs*_ and (1 − *m*_*s*_)*ρ*_*hs*_, and the term (1 − *m*)^2^*ρ*_*hs*_ by (1 − *m*)(1 − *m*_*s*_)*ρ*_*hs*_. We can check that the following qualitative properties holds true with a different mortality rate. However, it will modify the quantitative outcome of the model.

For any given pair of interaction traits, we can compute the equilibrium of this dynamical system.

#### Extinction

The extinction equilibrium, which corresponds to *ρ*_*ha*_ = *ρ*_*hs*_ = 0, always exists but it is unstable if

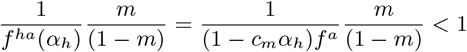

We have picked parameters, which fulfill this criterion (Table 1). In particular, we can see from this formula that increasing the mutualism cost *c*_*m*_ can lead to non viability of more mutualistic host. In our simulations, we fix this value to *c*_*m*_ = 0.3.

#### Absence of symbionts

We first look at the case where the symbionts are absent, *ρ*_*hs*_ = 0. Then the equilibrium *ρ*_*ha*_ = *ρ*_*h*_ should satisfies the following equation

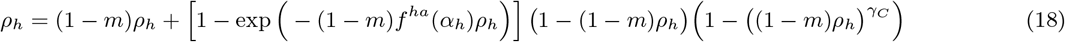

For our fixed parameters stated in Table 1, this equation admits a unique solution in [0, 1] and the proportion of host *ρ*_*h*_ alone without symbionts ranges between 0.25 and 0.34.

Without symbiont, the proportion of host converges to *ρ*_*h*_ = *ρ*_*ha*_ = 34, 14% (*γ*_*C*_ = 0.2 and *α*_*h*_ = 0). Thus in absence of any symbionts, the host always survives.

However, this equilibrium is unstable in our parameters range – the jacobian around this equilibrium has an eigenvalue *λ* with modulus greater than 1, *λ* = (1 − *m*)^2^ (1 + *ρ*_*h*_*f*^*s*^(*α*_*h*_, *α*_*s*_)). This suggests that a third equilibrium exists and may be stable.

#### Coexistence of symbionts and host

We also have a unique coexistence equilibrium which is a stationary state of the model (17). In our parameters range, this equilibrium always exists and it is always stable and attractive for any values of the interactions traits (*α*_*h*_, *α*_*s*_).

So, in the presence of a parasitic symbiont (*α*_*s*_ = 0), the proportion of hosts converges to *ρ*_*h*_ = 0.15 and the proportion of symbionts to *ρ*_*hs*_ = 0.106% which is in accordance with our simulation at initial time *t* = 0 (see Figure 2 b and c).

In addition, when hosts (*α*_*h*_ = 0) are associated with mutualistic symbionts (*α*_*s*_ = 1), the proportion of hosts rises to *ρ*_*h*_ = 0.638 and the proportion of symbionts to *ρ*_*hs*_ = 0.596. Thus the gain of cohabiting with mutualistic symbionts is indeed huge.

### A.3 Competition strength, perturbation and mutualism persistence

#### Competition strength determines mutualism persistence

In the main text, we show that competition is essential for the transition to mutualism, but it is also important for its persistence, as shown here. In this section, we explored the effect of a sudden variation in competition strength *γ*_*C*_ on the persistence of mutualism. We started with a strong competition *γ*_*C*_ = 0.2. As expected from our previous results, a transition to mutualism occurred (Figure A9a), c) and d)). Then, around *t* = 10^4^, we suddenly switched the competition strength to *γ*_*C*_ = 2, corresponding to negligible competition. We observed reversal of mutualism due to the proportion of mutualistic symbionts decreasing from 20% to less than 5% (Figure A9a) and e)). We observed that the reversal of mutualism due to a weakening of the competition corresponded with an increase in host and symbiont densities. This increase is due to the reduction of competition, which determines densities more than the presence or absence of mutualism does.

#### Density perturbation does not affect mutualism persistence

Next, we tested how mutualism responds to a decrease in competition due to eradication of hosts and symbionts in a large homogeneous region of space (Figure A10). While previously we demonstrated that mutualism regresses when competition is set to be weak, we show here that mutualism persists in the face of decreased competition due to decreased host density.

**Figure A8:**
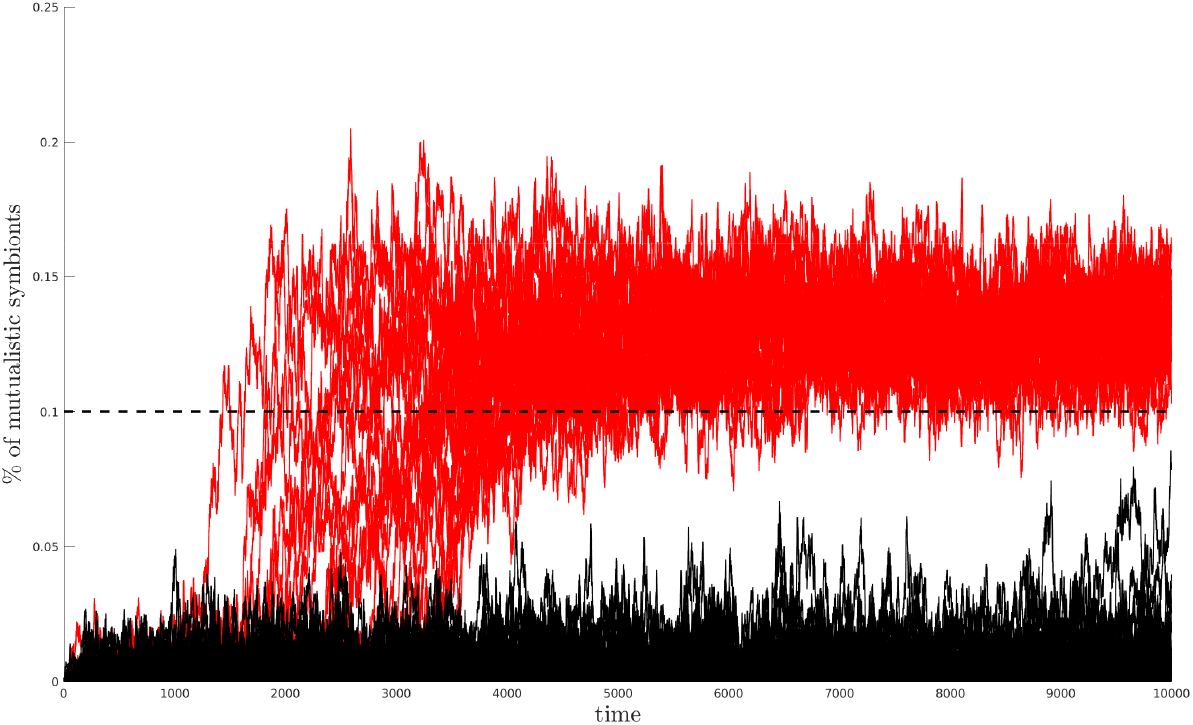
Evolution of the percentage of mutualistic symbionts in the population over 129 simulations. Red curves corresponds to replicates such that the percentage of mutualistic symbionts remains greater than the threshold of 10% – transition to mutualism. Black curves corresponds to replicates where the percentage remains below the 10% threshold – no transition.

**Figure A9:**
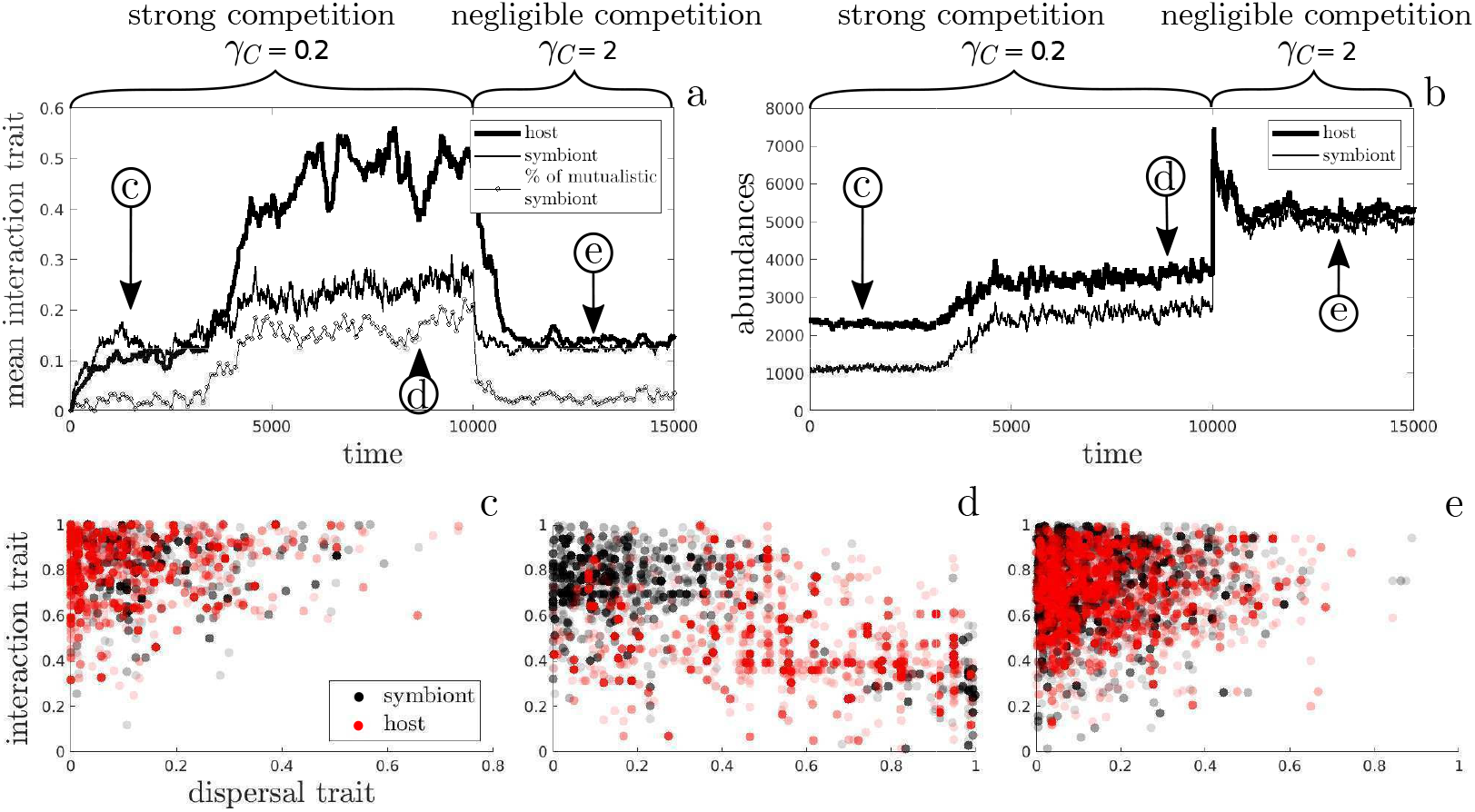
The transition to mutualism and collapse of mutualism depending on competition. a) Host and symbiont average interaction traits and the percentage of mutualistic symbionts over time. b) Host and symbiont abundance. There is strong competition from time *t* = 0 to *t* = 10^4^ and negligible competition from *t* = 10^4^ to the end. c) Dispersal and interaction traits distribution before the transition to mutualism. d) Dispersal and interaction traits distribution during mutualism persistence. e) Dispersal and interaction traits distribution after mutualism collapse. Parameters are 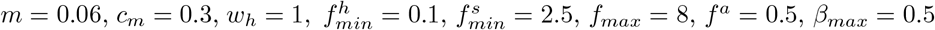 and *d* = 0.

### A.4 Mortality and dispersal cost can induce host dependency in emerging mutualistic systems

In the main text, we focused on the effect of mortality *m* and dispersal cost *d* on the transition to mutualism and host dependency. Here, we present in more detail the effect of dispersal cost on the distribution of hosts and symbionts in trait space for three values of dispersal cost and fixed mortality rate (Figure A11). In addition, the table A1 shows the features of the clusters in the trait distribution.

We demonstrated that the dispersal cost favours the transition to mutualism. Moreover, even when the cost was high, the features of the clusters revealed that parasitic symbionts maintained a more global dispersal than mutualistic symbionts.

**Figure A10:**
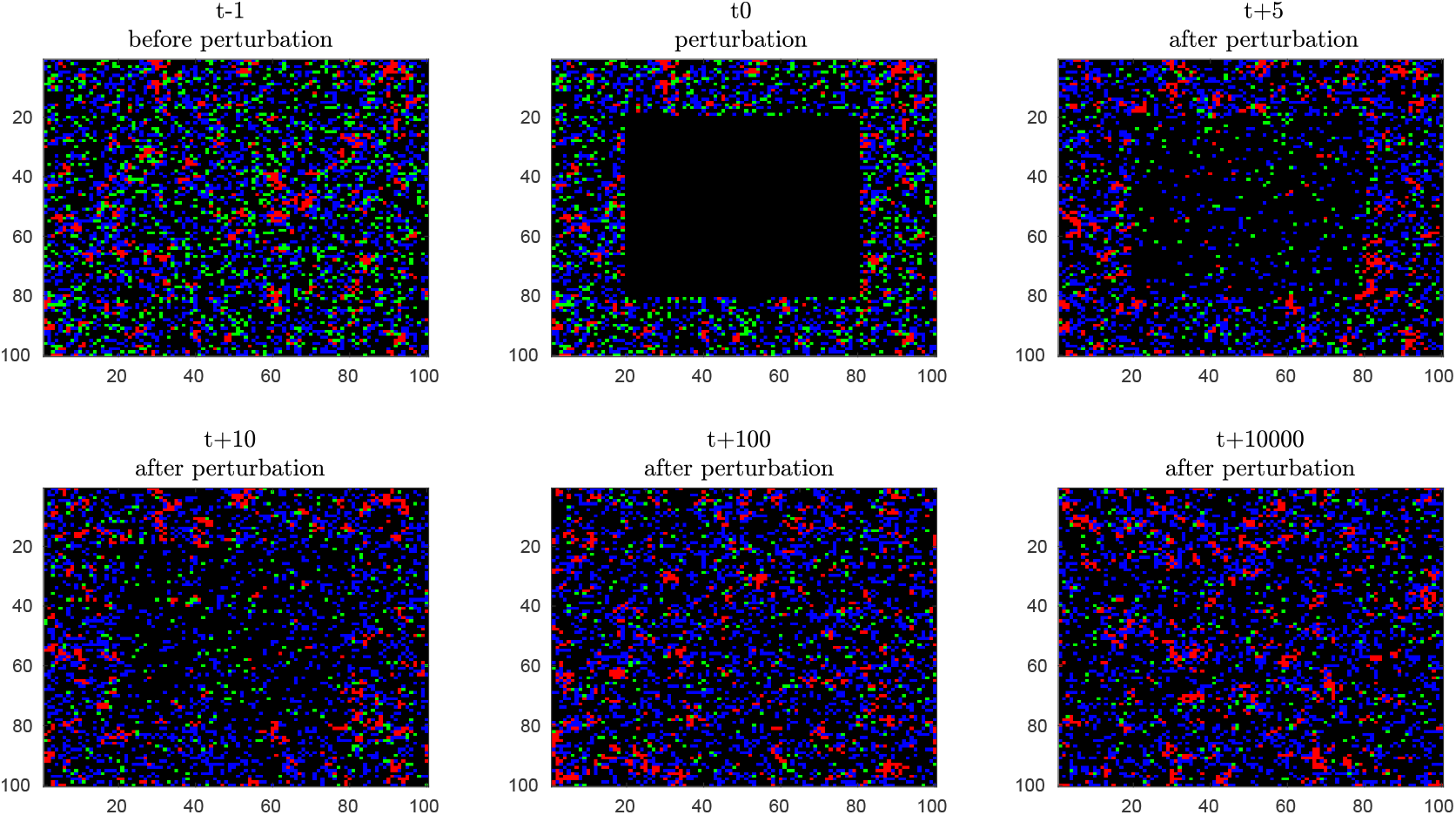
Maintenance of mutualism in the face of a reduction in competition caused by a perturbation creating a large square of free cells. Snapshot at several times: *t* − 1 is the eco-evolutionary equilibrium with mutualism just before the perturbation, then at *t*_0_ the perturbation, and then *t* + 5, *t* + 10, *t* + 100 and *t* + 10000 after the perturbation. In black, the free cells; in green, the hosts alone; in blue, the couples with parasitic symbionts; and in red, the couples with mutualistic symbionts. Parameters are 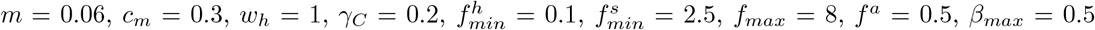 and *d* = 0.

**Figure A11:**
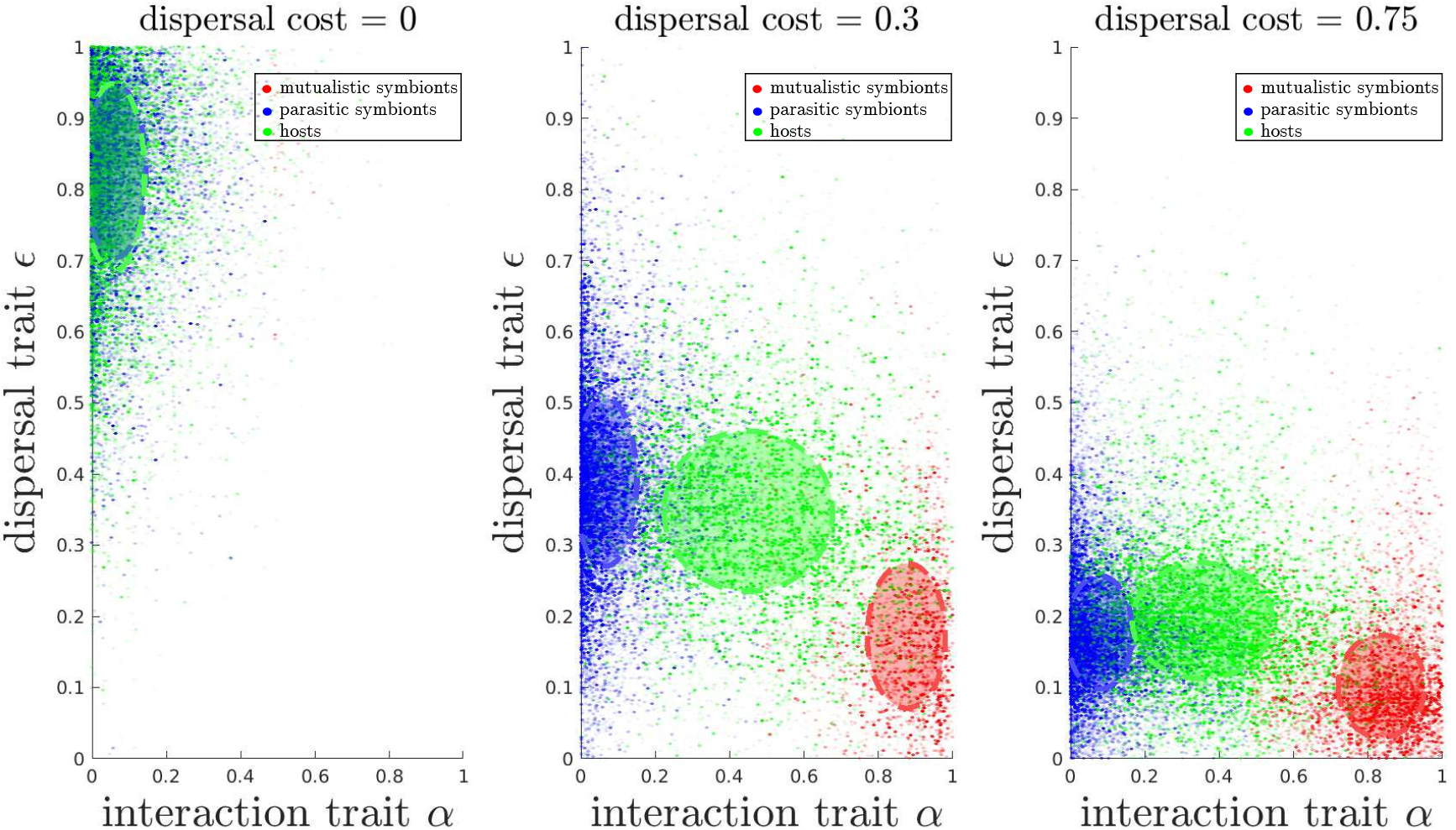
Joint distribution of the host (green), the parasitic symbiont (blue) and the mutualistic symbiont (red) populations in the traits domain for mortality *m* = 0.03 and an increasing dispersal cost *d* ∈ {0, 0.3, 0.75}. The ellipses correspond to the standard deviation. The 48 runs averaged in Figure 6 are plotted together. Other parameters are 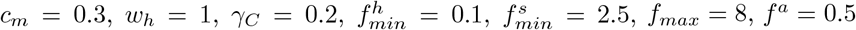 and *β*_*max*_ = 0.5.

**Table A1:**
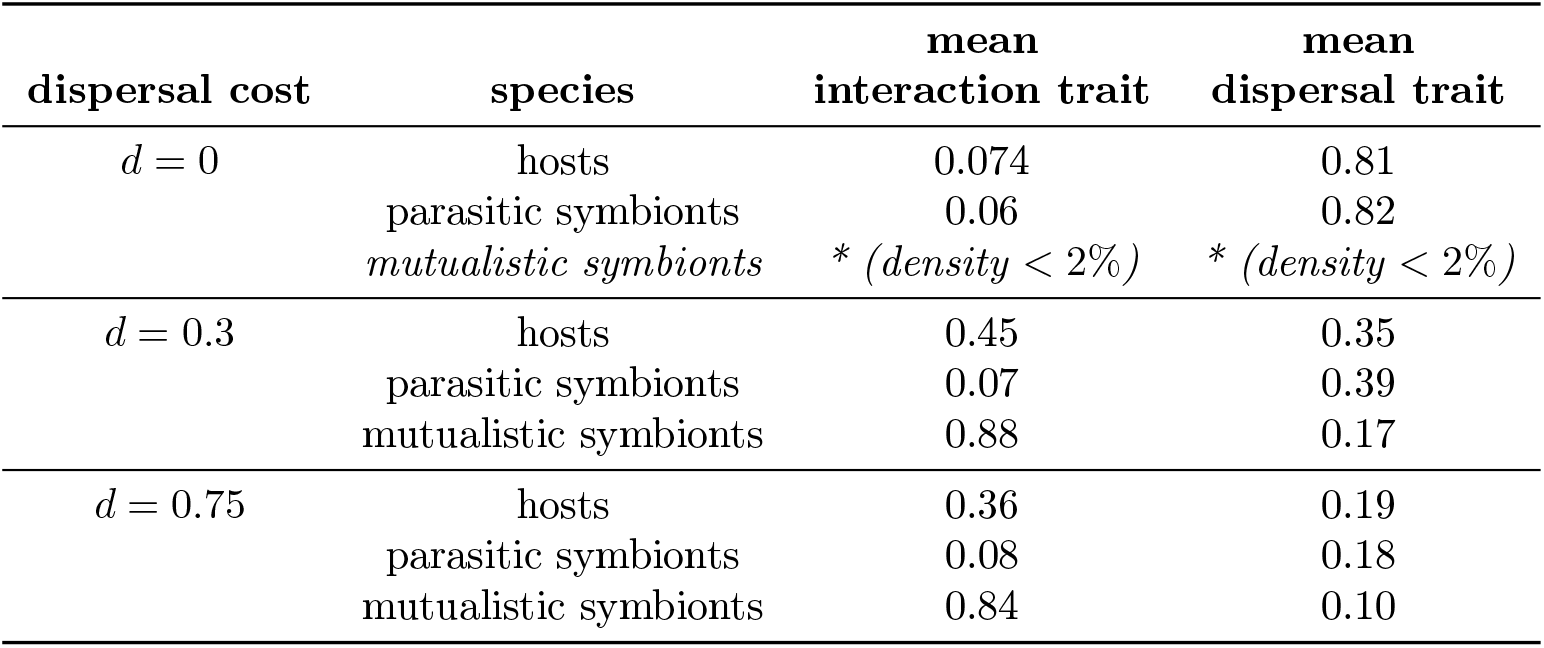
Features of the clusters in the traits domain

| dispersal cost | species | mean<br>interaction trait | mean<br>dispersal trait |
| --- | --- | --- | --- |
| $d = 0$ | hosts | 0.074 | 0.81 |
|  | parasitic symbionts | 0.06 | 0.82 |
|  | *mutualistic symbionts* | * (*density* < 2%) | * (*density* < 2%) |
| $d = 0.3$ | hosts | 0.45 | 0.35 |
|  | parasitic symbionts | 0.07 | 0.39 |
|  | mutualistic symbionts | 0.88 | 0.17 |
| $d = 0.75$ | hosts | 0.36 | 0.19 |
|  | parasitic symbionts | 0.08 | 0.18 |
|  | mutualistic symbionts | 0.84 | 0.10 |

### A.5 Density-dependent competition between symbionts

In the main text, symbionts compete for free hosts, which is a form of density-dependent competition. Other ecological factors may also lead to density-dependent competition between symbionts, for instance if symbionts compete for resources that are not provided by the hosts. Figure A12b shows that density-dependent competition between symbionts reduces symbiont density, as expected. Hosts are therefore free of symbionts more often, which selects for non-mutualistic hosts (Figure A12c, to be compared with Figure 3b).

**Figure A12:**
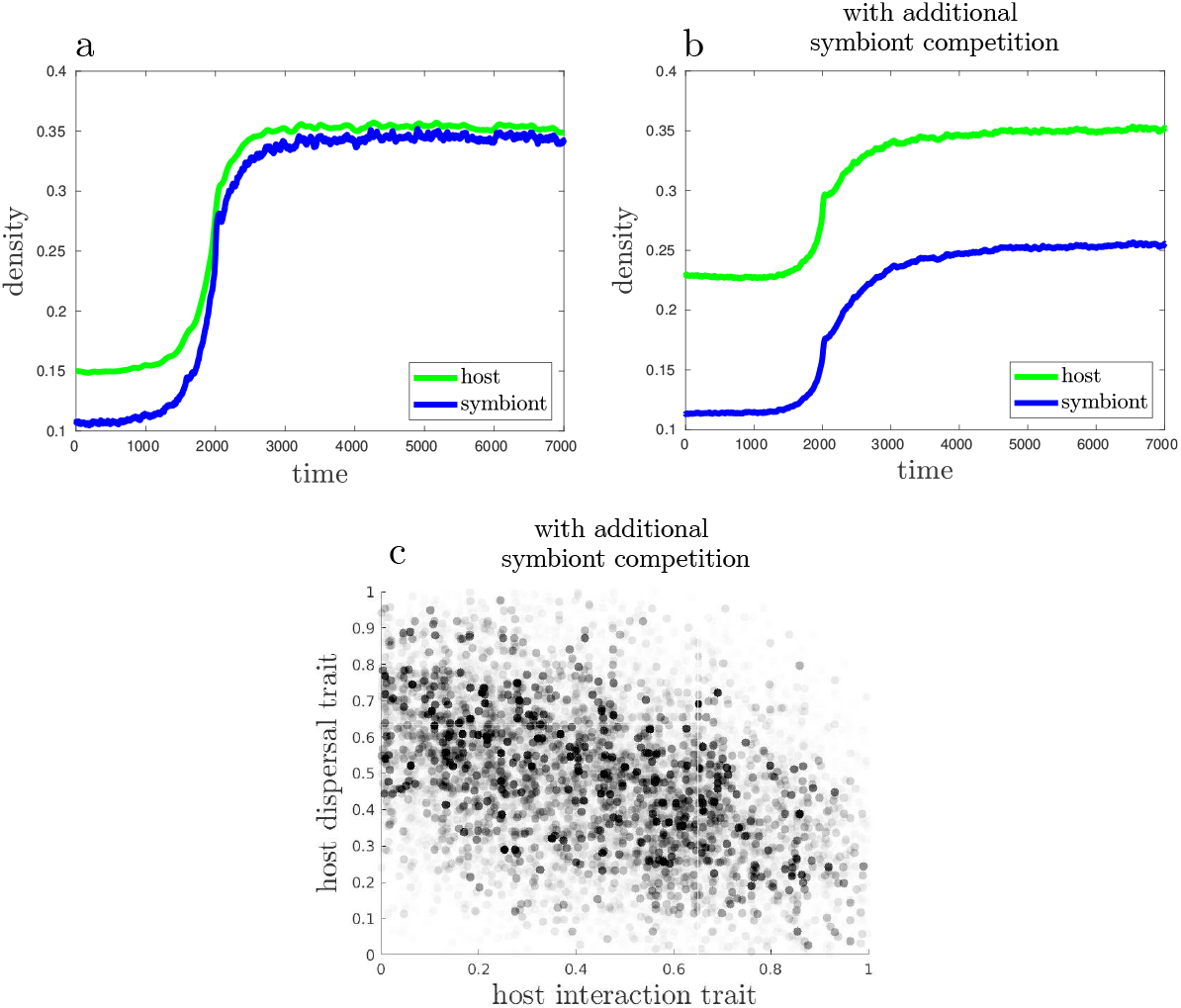
Effect of density-dependent competition between symbionts on host evolution. a) Symbiont and host densities after and before the transition, without density-dependent competition between symbionts. b) Symbiont and host densities after and before the transition, with density-dependent competition between symbionts. c) Distribution of host traits after the transition, with density-dependent competition between symbionts. To be compared with Figure 3b.

### A.6 Evolutionary rescue

Figure 6 provides evidence for evolutionary rescue, as discussed in the main text. Figure A13 shows that this occurs only in a fraction of the simulations, when mutualists arise soon enough to rescue the whole system.

**Figure A13:**
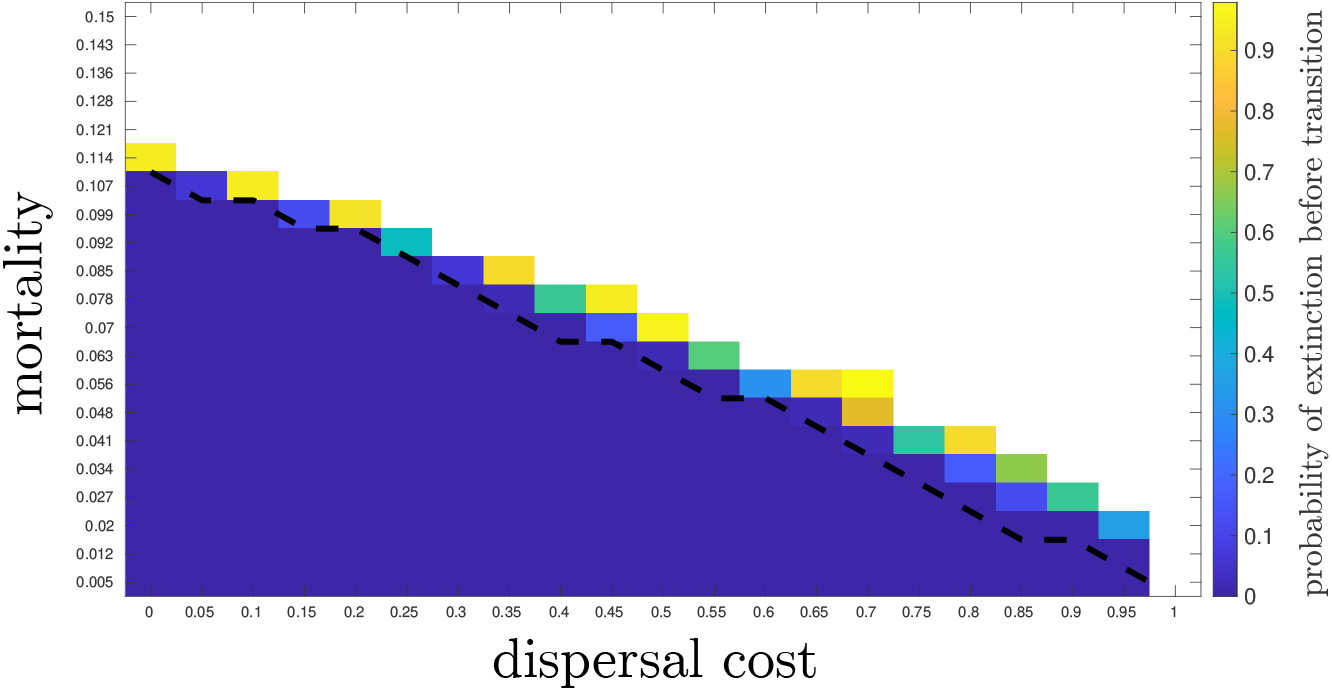
Frequency at which evolutionary rescue occurs. This figure is identical to Figure 6, except that it maps the probability of extinction before the transition, instead of the mean percentage of mutualistic symbionts. The dotted black line indicates the upper boundary of viability for the parasitic system, without evolution. Above the dotted black line, in some cases the evolution of mutualism rescued the whole system, although the parasitic system is unviable alone. In the white region, the systems goes extinct, even with evolution

### A.7 Symbionts competition within hosts

In our current model, a host can be colonised only by one symbiont and once the symbiont is established on a host, it cannot be replaced by another symbiont. Furthermore, when several symbionts arrive at the same time on an available host, the symbiont, which establishes, is chosen randomly uniformly among the contenders. Here, we relax these assumptions in order to model symbionts’ competition within a host, or “superinfection”. We assume that within a host, the most parasitic symbiont, with the lowest interaction trait, is the most competitive symbiont. Thus, it will be more efficient to establish in a host or dislodge a symbionts from the host.

#### Establishment of symbionts on a host

Specifically, when *N* symbionts, with trait {*α*_1_, …, *α*_*N*_ } arrive on a host, the establishment probability 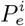 of the symbiont *i* is given by :

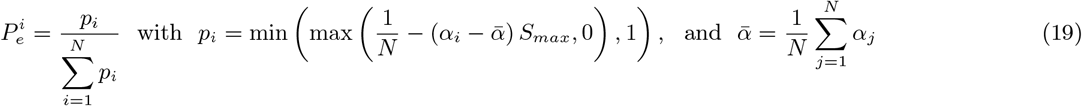

where *S*_*max*_ measures the superinfections’ intensity, which corresponds to the maximal competitive advantage of a symbiont. For instance, when a truly parasitic symbiont *α*_1_ = 0 tries to establish with a truly mutualsistic symbiont *α*_2_ = 1, its establishment probability is 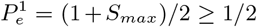. The establishment probability of the mutualsitic symbiont is 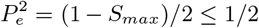. If *S*_*max*_ = 0, they have the same probability of establishment, while if *S*_*max*_ = 1, the parasitic symbiont always over-competes the mutualistic symbiont.

#### Replacement of a resident symbiont

When *N* symbionts with trait { *α*_1_, …, *α*_*N* }_ arrive in a host already occupied by a resident symbiont with trait *α*_*s*_, they may dislodge the resident. Specifically, the probability of the resident symbiont to persist *P*_*p*_ is given by

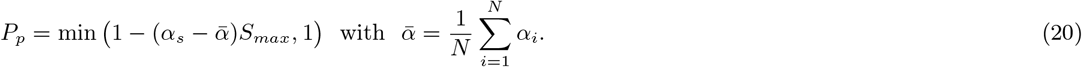

In particular, if the resident has a trait *α*_*s*_ lower than the mean trait of the invaders 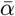, then the resident always persists. Otherwise, the resident may be dislodged with a probability smaller than *S*_*max*_. Then if the resident is dislodged, the establishment probability of the *N* invader symbionts is given by the previous formula (19).

Figure A14 shows the effect of the superinfection intensity *S*_*max*_ on the percentage of mutualistic symbionts. We show that despite the competitive advantage of parasitic symbionts when competing for a host, the transition to mutualism is possible when the superinfection intensity is not too large (if *S*_*max*_ *<* 1*/*2, transition occurs, that is the percentage of mutualistic symbionts stays above 10%). Moreover, when *S*_*max*_ *<* 1*/*2, the trait distribution of symbionts is bimodal.

**Figure A14:**
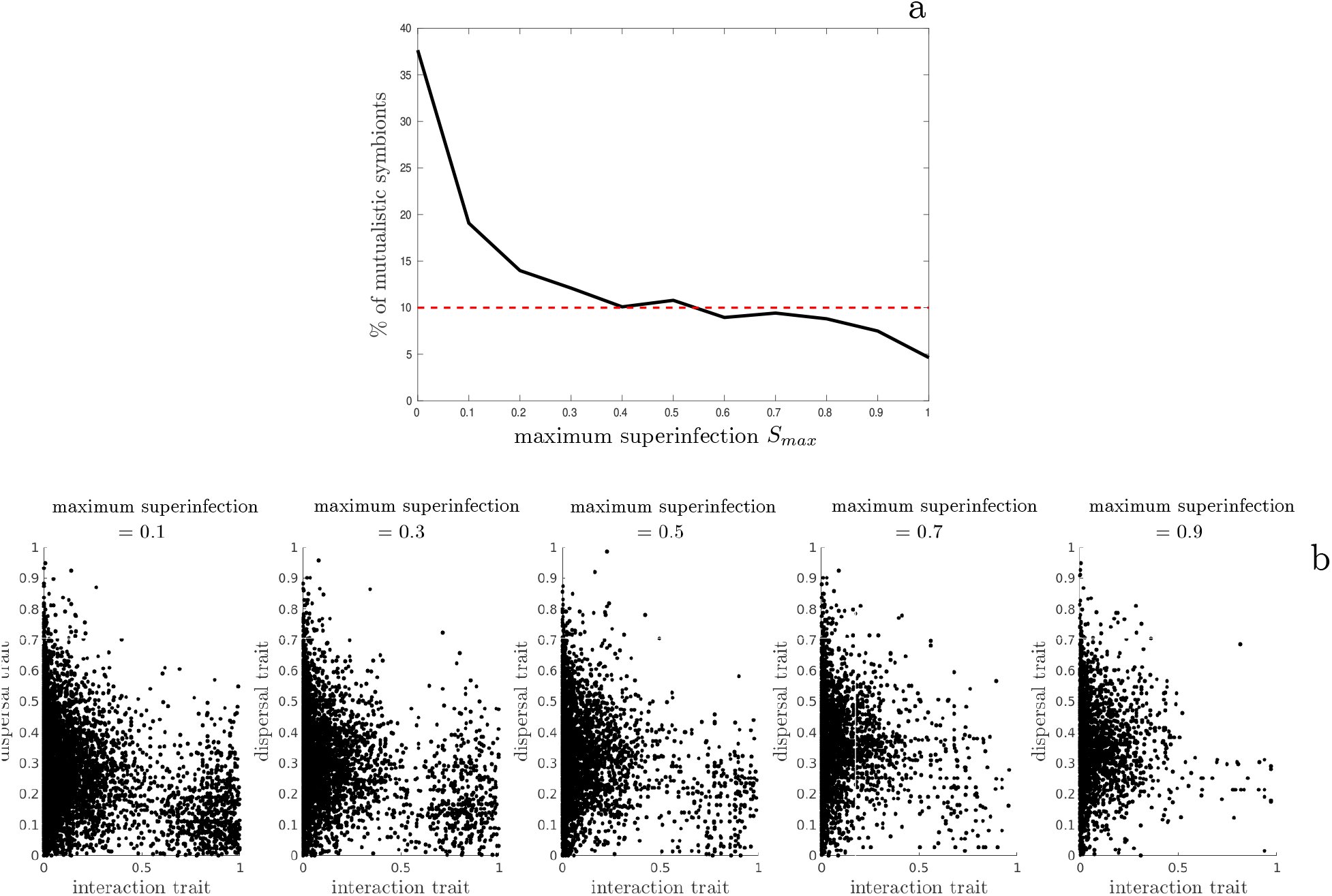
a) Percentage of mutualistic symbionts in function of the maximum superinfection advantage *S*_*max*_ averaged over 20 simulations per parameter values. b) Distributions of symbionts population in traits domain according to five intensity of superinfection advantage *S*_*max*_. Distributions corresponds to 20 simulations for each parameter values. These results are obtained with a maximum time projection of 5000 time steps, a strong and global competition (*γ*_*C*_ = 0.2) and a dispersal cost (*d* = 0). Others parameters are 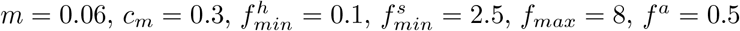 and *β*_*max*_ = 0.5.

## Footnotes

1 Symbiosis is used here in its etymological sense of “living together”, encompassing parasitic and mutualistic symbiosis.

2 An altruistic trait benefits conspecifics, at a cost to its bearer. In contrast, a mutualistic trait benefits heterospecifics.

